# The axonal actin-spectrin lattice acts as a tension buffering shock absorber

**DOI:** 10.1101/510560

**Authors:** Sushil Dubey, Nishita Bhembre, Shivani Bodas, Aurnab Ghose, Andrew Callan-Jones, Pramod A Pullarkat

## Abstract

Axons are thin tubular extensions generated by neuronal cells to transmit signals across long distances. In the peripheral and the central nervous systems, axons experience large deformations during normal activity or as a result of injury. Yet, axon biomechanics, and its relation to the internal structure that allows axons to withstand such deformations, is poorly understood. Up to now, it has been generally assumed that microtubules and their associated proteins are the major load-bearing elements in axons. We revise this view point by combining mechanical measurements using a custom developed force apparatus with biochemical or genetic modifications to the axonal cytoskeleton, revealing an unexpected role played by the actin-spectrin skeleton. For this, we first demonstrate that axons exhibit a reversible strain-softening response, where its steady state elastic modulus decreases with increasing strain. We then explore the contributions from the various cytoskeletal components of the axon, and show that the recently discovered membrane-associated skeleton consisting of periodically spaced actin filaments interconnected by spectrin tetramers play a prominent mechanical role. Finally, using a theoretical model we argue that the actin-spectrin skeleton act as an axonal tension buffer by reversibly unfolding repeat domains of the spectrin tetramers to buffer excess mechanical stress.

## INTRODUCTION

Axons are micron-thin tubular extensions generated by neuronal cells to transmit signals across large distances — up to the order of centimeters in the brain and up to a meter in the peripheral nervous system of a human body [1]. In order to achieve such extreme aspect ratios, axons have evolved a unique organisation of the cytoskeleton. It has an axisymmetric structure with a central core of aligned microtubules arranged in a polar fashion and cross-linked by associated proteins (Fig. 1). This core is surrounded by neurofilaments and a membrane-associated cortex of cross-linked actin filaments [1]. Recent experiments using super-resolution light microscopy have revealed that the actin cortex also involves a periodic array of F-actin rings which are interconnected by spectrin cross-bridges [2, 3]. A myriad of proteins, including motor proteins, interconnect these structural elements to form a dynamic composite gel.

**FIG. 1.**
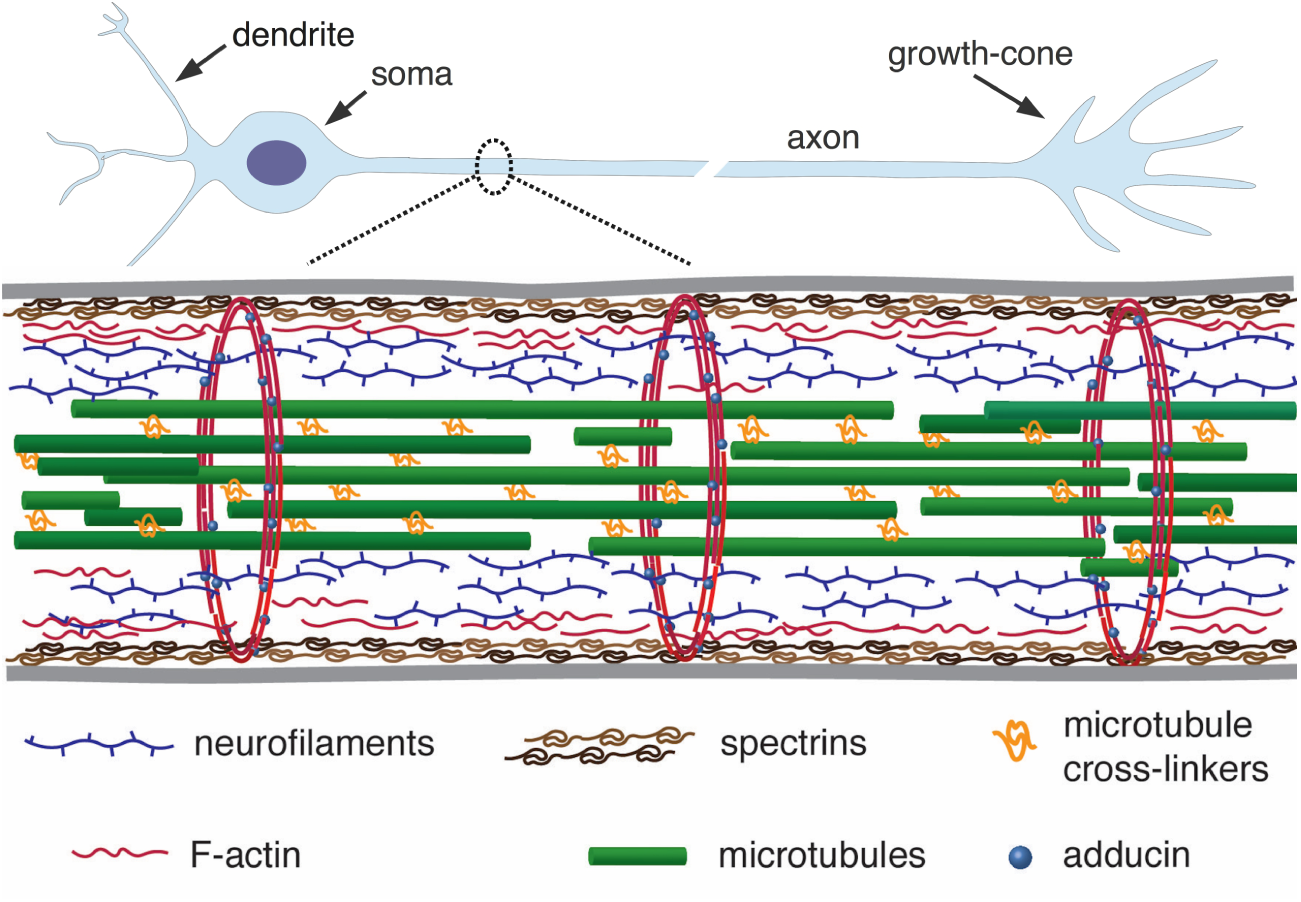
A simplified schematic of the axonal cytoskeleton. Typically, the axonal core has a bundle of microtubules which are cross-linked by a variety of microtubule associated proteins which includes tau (in some cases a more loose organisation of microtubules interdispesed with neurofilaments is seen) [17]. This core is surrounded by neurofilaments. The outer most scaffold has an array of periodically spaced rings composed of F-actin filaments. The actin rings are interconnected by *α*/*β*-spectrin tetramers, which are aligned along the axonal axis (only tetramers in a cross-section are shown for clarity). Other cortical F-actin structures also exist. A myriad of proteins, including motor proteins (not shown) interconnect the various filaments, and also the membrane (grey lines) to the inner skeleton. The chick DRG axons we use are about 1 µm thick and the rings in them are about 200 nm apart.

Axons can be subjected to large stretch deformations under a variety of normal as well as abnormal conditions. Mammalian sciatic nerves, for example, can experience localized strains up to 30% during regular limb movements [4]. A much more extreme case of stretching occurs in the nerves running along the mouth-floor of certain species of baleen whales where it can reach up to 160% during feeding, and these axons have evolved special strategies to cope with such situations [5]. Recent *in-vivo* studies in humans have shown significant changes in nerve stiffness due to limb flexion [6]. This suggest that nerve tissue may possess unique mechanical behavior, perhaps to offer some protection against stretch injuries, which are common. It is also known that the brain, being one of the softest tissues, undergoes significant shear deformations (2–5% strain) even under normal activities such as jumping [7]. At higher accelerations, transient to traumatic brain concussion occur and is a leading cause of injury in contact sports, with an estimated 1.6 to 3.6 million cases annually in the USA alone [8]. Hence, knowing how and under what conditions brain deformation compromises axonal integrity is considered essential to understand, manage, and treat concussions [9, 10].

At the single cell level, several studies have explored axonal responses to stretch deformations, primarily using glass micro-needles to pull on live axons and video microscopy to record the evolution of force and strain. Pioneering experiments by Heidemann and co-workers show that axons respond viscoelastically to stretch deformations [11]. Besides, they have shown that axons can also develop excess tension when stretched, presumably due to the action of molecular motors [11]. Such contractile stress generation has since been studied experimentally and theoretically in some detail and is found to arise due to the action of myosin-II molecular motors [12–15]. Despite the observation of such a rich variety of mechanical responses, the structure–function relationships between the different cytoskeletal elements and these mechanical responses are poorly understood.

In this article, we use a strain-controlled, optical fiber based force apparatus to show that axons exhibit a hitherto unknown strain-softening behavior where the steady state elastic modulus diminishes with increasing strain. By combining our extension rheology technique with biochemical and genetic interventions that alter specific cytoskeletal components, we demonstrate that the actin-spectrin skeleton is a major contributor to the axonal mechanical response to stretch. Then, with the help of a theoretical model, we argue that strain-softening possibly arises from the force-induced unfolding of spectrin subunits, a process known to occur when single spectrin molecules are stretched [16]. Based on these findings we propose that the actin-spectrin skeleton in axons can act as a tension buffer or “shock-absorber” allowing axons to undergo significant deformations without excess build up of tension, as seen in our experiments. Moreover, we show that this mechanism renders axons with a viscoelastic solid-like response, with memory of the initial state, which allow axons to undergo reversible stretch deformations.

## RESULTS

### Axons exhibit strain-softening, viscoelastic solid-like response

In order to study the axonal response as a function of imposed strain, we applied successively increasing strain steps with a wait time between steps using the home-developed force apparatus shown in Fig. 2A (image of setup: Fig. S1, stretching of axon: Movie-S1) [18]. After each step, the strain is held constant using a feedback algorithm. The resulting force relaxation data is shown in Fig. 2B. Unless specified otherwise, all data are for cells grown for two days *in-vitro* (2-DIV). The force relaxation after each step is indicative of the viscoelastic nature of the axon. From the data, we can calculate the axonal tension 𝒯 = *F*/(2 sin *θ*) (see Fig. 2C), where 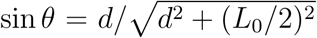, with *θ*(*t*) as the angle with respect to the initial position, *d* is the displacement of the tip of the cantilever which is in contact with the axon mid point, and *L*_0_ is the initial length of the axon. The calculated tension plotted in Fig. 2B shows that it tends to relax to the same steady state value 𝒯_ss_ as the strain on the axon increases. The inset of Fig. 2D shows this trend for 𝒯_ss_ seen for multiple axons. The occurrence of a steady state tension (or a steady state force) is indicative of a solid-like behavir of the axons at longer timescales. To check this further we performed a few experiments with wait time ∼ 10 min ≫ the typical relaxation time (quantified later) after a strain step and these show that the force indeed decays to a non-zero steady state value (Fig. S2).

**FIG. 2.**
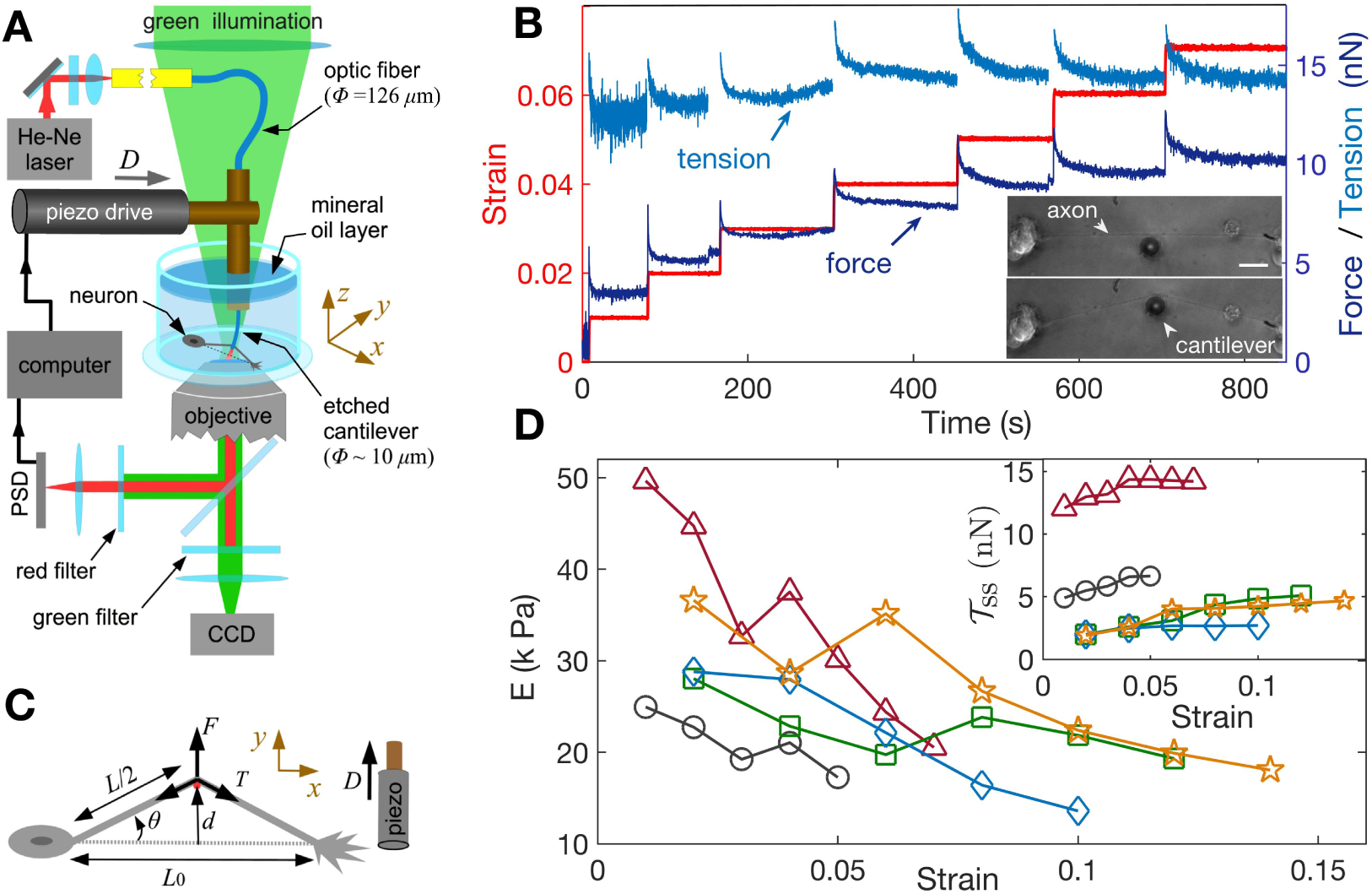
Feedback-controlled axon stretch apparatus reveals strain-softening behavior. **(A)** The schematic of the home-developed force apparatus that uses an etched optical fiber as a cantilever to stretch axons and to sense force. Laser light exiting the cantilever tip is imaged on to a Position Sensitive Detector (PSD), which in turn is read by a computer. The computer controls a piezo drive via a feedback algorithm to apply strain steps and then to maintain the strain constant after each step. **(B)** Typical force response (dark blue) of a 2-DIV axon to increasing strain explored using successive strain steps (red). The calculated tension in the axons (sky blue) is also shown. The inset shows the images of the axon before and after stretch (scale bar: 20 µm). The light exiting the etched optical fiber cantilever can be seen as a bright spot (reduced in intensity for clarity), and this is imaged on to the PSD to detect cantilever deflection. **(C)** Illustration of the parameters used in the calculations. The strain is calculated as 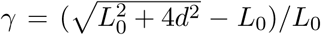, and force on the cantilever as *F* (*t*) = −*k*(*D* − *d*), where *k* is the cantilever force constant. The axonal tension is 𝒯 = *F*/(2 sin *θ*), where 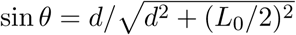. **(D)** Young’s moduli calculated for different 2-DIV axons using the steady state tension after each step show a strain-softening behavior (the different symbols are for different axons). Only a few representative plots are shown for clarity, and more examples are shown in Fig. S4. The inset shows the tension vs. strain plots for different axons. Tension tends to saturate with increasing strain (tension homeostasis), which leads to the observed softening.

We note that the axonal structure is highly anisotropic and is composed of multiple cytoskeletal structures, and its elastic response is non-linear. Considering extensile deformations alone, we calculate an “effective” Young’s modulus *E* = (𝒯_ss_ − 𝒯_0_)/(*A.γ*), where 𝒯_0_ is the tension of the axon at zero strain which is calculated by extrapolating the tension vs. strain data (Fig. S3), *A* is the cross-sectional area of the axon (neglecting the change in *A* with strain), and 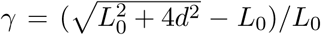 is the imposed strain. This modulus is expected to be different from that measured using AFM or magnetic tweezers where the imposed force or strain is radial [19, 20]. The effective modulus data we obtained from multiple axons is shown in Fig. 2D, with more examples in Fig. S4. Remarkably, the elastic modulus shows a strain-softening which actually reflect the tendency for tension to saturate with increasing strain (tension homeostasis). This strain-softening response is not due to any permanent damage or plastic flow as the response remains qualitatively the same when repeated for the same axon. Moreover, when the axons are released from the cantilever from the maximal strain state they recover their initial length within 5–10 s (Movie-S2).

### F-actin plays a more significant role than microtubules in axonal response to stretch

Since turnover of cytoskeletal elements, such as actin and microtubules, can be strain-dependent, for example, via opening of stretch-activated *Ca*^++^ channels, it is conceivable that the strain-softening response arises due to cytoskeletal remodelling. To test this, and the relative contributions of different structural elements, we performed experiments aimed at either stabilizing or destabilizing microtubules and F-actin using 2-DIV axons. We first tested the response of axons treated with the microtubule stabilizer Taxol at 10 µM for 30 min and the resulting data are shown in Fig. 3A. The elastic moduli show a strain-softening response similar to normal axons, albeit with higher values of the moduli. Next, in order to check whether F-actin dynamics could lead to strain-softening, we treated neurons with 5 µM of the F-actin stabilizing agent Jasplakinolide for 30 min; see Fig. 3B. As shown in Fig. 3C, F-actin-stabilized axons exhibit a more substantial increase in the elastic moduli compared to Taxol treatment, while both display pronounced strain-softening.

**FIG. 3.**
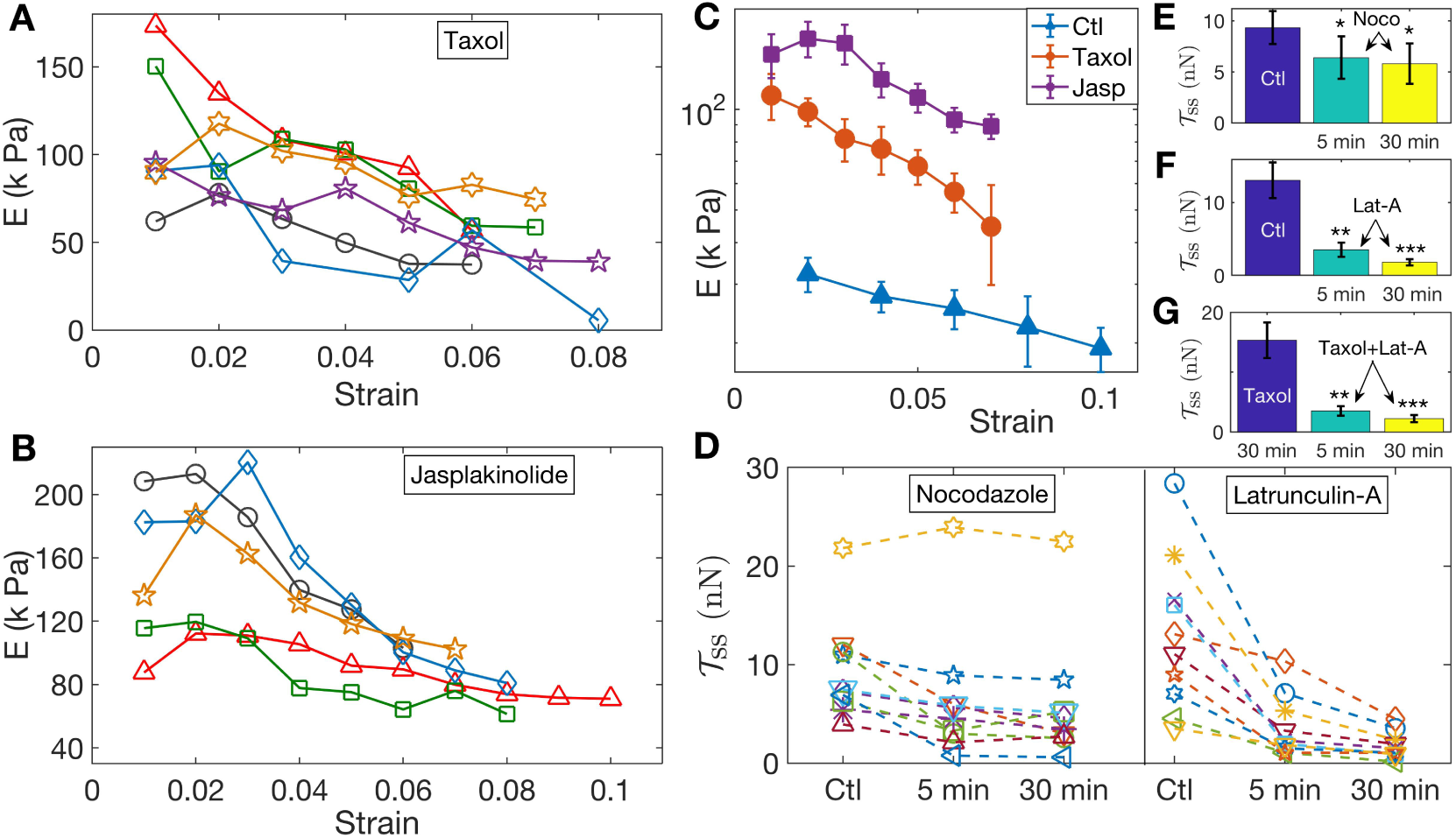
Pharmacological perturbations of the cytoskeleton reveals a key role for F-actin in the axonal stretch response. **(A,B)** Stabilizing microtubules or F-actin by treating 2-DIV axons with either 10 µM Taxol or 5 µM Jasplakinolide leads to significant increases in axonal stiffness, but the strain-softening response persists (more data in Figs. S5, S6; the different symbols are for different axons). **(C)** Log-linear plots of the averaged elastic moduli as a function of strain for different treatments as well as control (for data shown in A,B and Fig. 2D). It can be seen that stabilization of F-actin causes a larger increase in moduli compared to stabilizing microtubules (error bars are standard error (SE)). **(D)** Change in steady state tension 𝒯_ss_ obtained after treating 2-DIV neurons with either the microtubule disrupting drug Nocodazole (10 µM) or the F-actin disrupting drug Latrunculin-A (1 µM) (n = 10 each). These treatments leave the axons very fragile and they detach easily when pulled. Hence a cyclic step-strain protocol with *γ* = 0.01 was employed as detailed in the text. As 𝒯_0_ cannot be determined from single steps, we compare the net tension for the same axon before (Ctl) and after 5 min and 30 min of treatment. The data show a significant decrease in axonal tension after F-actin disruption and a relatively smaller decrease after disrupting microtubules. **(E, F)** Bar plots of 𝒯_ss_ showing the significance of the two treatments shown in D. Tension was measured for the same axons 5 min and 30 min after exposing to drug (error bars are SE). **(G)** Plots of 𝒯_ss_ for axons initially treated with Taxol for 30 min, and for the same axons subsequently treated with Lat-A for 5 min and 30 min (n = 7).

To further investigate the possible role of the microtubule and F-actin cytoskeletons, we performed axon stretch experiments after specifically disrupting each of these elements. To avoid irreversible damage to drug-treated, cytoskeleton-weakened axons, instead of the previous increasing step-strain experiments, we subjected them to a cyclic strain protocol where repeated up and down steps of equal magnitude are applied (Fig. S7). Moreover, to reduce scatter in data due to natural axon to axon variation, the same axon was probed before and after treatment. We first applied cyclic strain on control axons for extended periods to ensure that axons are not damaged under such conditions (Fig. S7). We then performed measurements after depolymerizing microtubules using Nocodazole (Noco) at 10 µM for up to 30 min. After this treatment, some axons exhibited pronounced beading as is expected when microtubules are lost. As can be seen from Fig. 3D, on the average, treated axons showed a reduction in steady state tension: 𝒯_ss_(*γ* = 0.01) = 9.3 nN ± 1.6(control), 6.4 nN ± 2.1(Noco: 5 min), 5.8 nN ± 2.0(Noco: 30 min), where the values are mean ± SE (see Fig. 3E, and individual data in Fig. 3D). Next, we disrupted F-actin using 1 µM Latrunculin-A (Lat-A) for up to 30 min and observed that this treatment produced a much more dramatic reduction in the steady state tension than did Nocodazole, as shown in Fig. 3D. In this case, we obtained 𝒯_ss_(*γ* = 0.01) = 13.0 nN ± 2.5(control), 3.5 nN ± 1.0(Lat-A: 5 min), 1.8 nN±0.4(Lat-A: 30 min), where the values are mean ± SE (see Fig. 3F, and individual data in Fig. 3D). This correlates well with the tendency of Jasplakinolide to cause a larger increase in axon elastic modulus as compared with Taxol (see Fig. 3C). In addition, the tension relaxation is much faster for Latrunculin-A treatment as compared with either Nocodazole-treated or control axons (Figs. S8, S9), suggesting that F-actin plays a leading role in strain-softening in axons.

The larger effects seen after F-actin stabilization or disruption, when compared to microtubule perturbation, comes as a surprise as axonal mechanics is thought to be dominated by microtubules. Since there are interdependencies between the stability of these two component, we tested whether the sharp decrease in moduli after F-actin disruption could be due to microtubules too becoming destabilized subsequent to Lat-A treatment. For this we first treated axons with Taxol to stabilize microtubules, and then exposed these axons to Lat-A (all concentrations and time as above). This data, presented in Fig. 3G, shows a drastic reduction in the steady state tension after Lat-A treatment of microtubule stabilized axons. The values are 𝒯_ss_(*γ* = 0.01) = 15.0 nN ± 3.0(Taxol), 3.5 nN ± 0.8(Taxol+Lat-A: 5 min), 2.2 nN ± 0.6(Taxol+Lat-A: 30 min).

The emergence of F-actin as more relevant than microtubules to the axon mechanical response under stretching is surprising given that microtubules usually form an aligned and tightly cross-linked bundle at the core of the axons. This finding prompted us to explore in more detail how F-actin regulates the axon stretch response, in particular via the periodic lattice of rings it forms together with *α* and *β* spectrin tetramers [2, 3].

### Spectrin contributes prominently to axonal stretch response

To check the spectrin distribution along axons we first imaged spectrin content in chick DRG axons using antibody labelling and confocal microscopy. Every axon imaged (n = 180) showed significant spectrin fluorescence distributed along the axon even at 2-DIV (Fig. S10). The ultra-structure of the spectrin organisation is revealed in the STED nanoscopy images shown in Figs. 4A1, 4A2, S11. The periodic lattice becomes more prevalent with the number of days the neurons are in culture while maintaining a periodicity in the range of 190 to 200 nm (see Table-S1 and Fig. S12 for quantification). The effect of the different drug treatments on the periodic lattice is summarised in section VII-D of Supplementary Material. Most notably, the periodicity is lost after Lat-A treatment (see Fig. S13 for quantification).

**FIG. 4.**
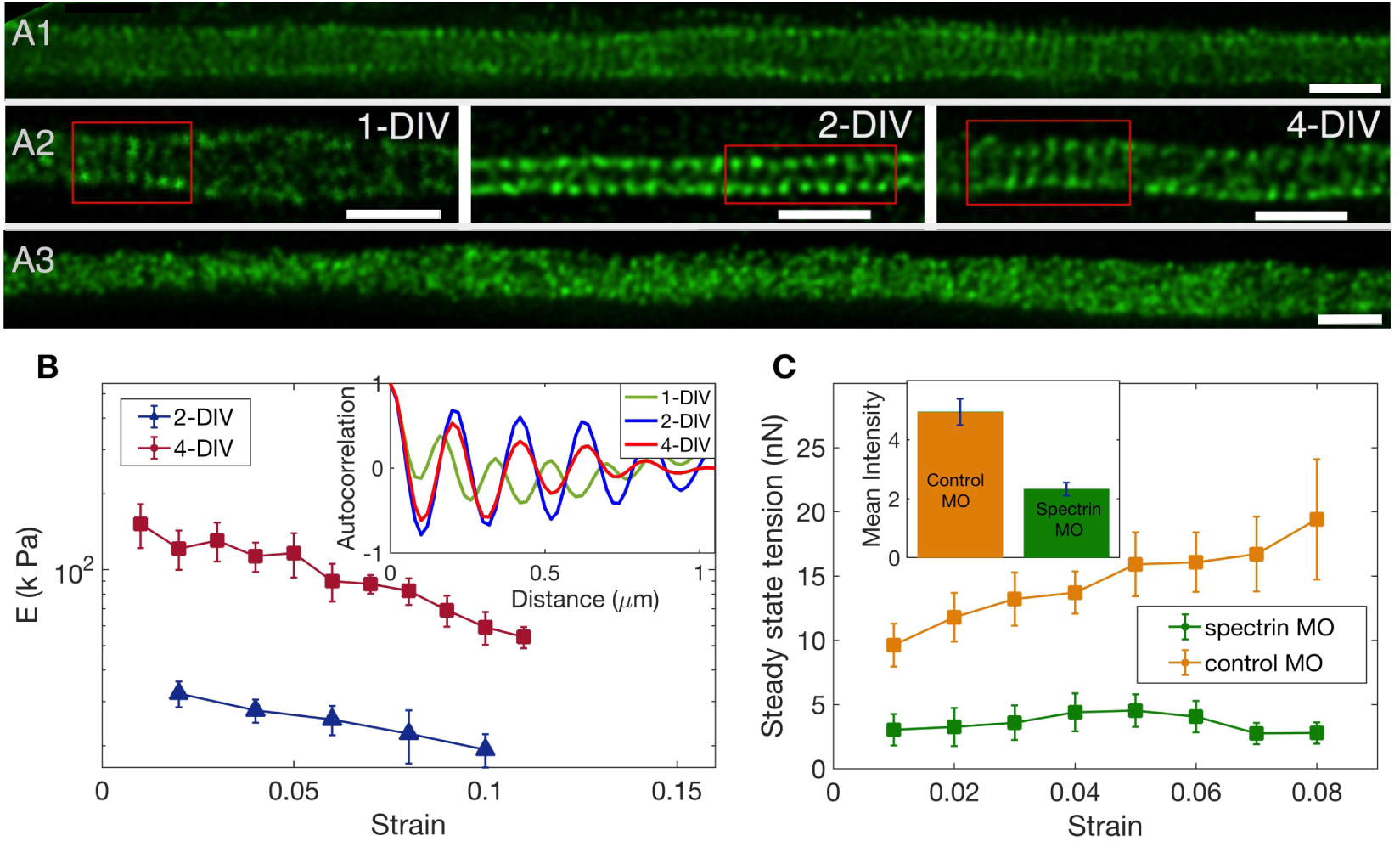
The role of spectrin in axonal mechanics. **(A1)** STED nanoscopy image of a 2-DIV axon immunolabelled with *β*-II spectrin antibody (vehicle control). This axon shows rings all along the imaged segment (see Fig. S11 for more examples). **(A2)** STED images of *β*-II spectrin in control axons of different DIVs. Observation of a large number of axons shows that the spectrin periodicity becomes more prevalent with age of the axon (see Table-S1). **(A3)** STED image of a 2-DIV axon treated with 1µM Lat-A for 30 min shows disordered *β*-II spectrin intensity distribution (see Fig. S13 for quantification). All scale bars are 1 µm. **(B)** Log-linear plots of averaged Young’s modulus vs strain for axons at 2-DIV (n = 5) and 4-DIV (n = 7) show a marked increase in moduli with age (error bars are SE). This increase in moduli correlated with the prevalence of the actin-spectrin lattice as a function of DIV which is quantified in Table-S1. The inset shows sample intensity-intensity autocorrelation functions for different DIV axons. The periodicity corresponds to the *α*/*β* spectrin tetramer length which is about 190 nm. **(C)** Plots of the averaged steady state tension 𝒯_ss_ as a function of strain for 2-DIV neurons treated with a non-specific, control morpholino (n = 10) and for those treated with an anti-*β*-II spectrin morpholino (n = 10). There is a substantial reduction in the tension measured after knock-down. The inset shows the extent of the knock-down obtained using antibody labelling of spectrin for control (n = 90) and knock-down neurons (n = 91); all error bars indicate the standard error (p < 0.05) (see Materials and Methods for details).

Remarkably, the increase in prevalence of the actin-spectrin lattice with age quantified in Table-S1 correlates well with the mechanical response, with axons of older neurons showing much higher values of Young’s moduli for all strain values as can be seen from Fig. 4B. The axonal rest tension, too, increases slightly with days in culture from 𝒯_0_(2-DIV) = 3.7 nN±1.2 (n = 10) to 𝒯_0_(4-DIV) = 5.3 nN ± 2.2 (n = 7), where the values are mean ± SE (Fig. S14). The variation of the steady state tension 𝒯_ss_ with strain for 4-DIV cells shows that tension tends to saturate, and is shown in Fig. S15.

Next, we performed knock-down experiments using a specific morpholino (MO) against chick *β*-II spectrin. Depletion of *β*-II spectrin has been reported to abolish the development of the periodic organization of the actin-spectrin membrane-associated skeleton [21]. Anti-*β*-II spectrin morpholino-treated axons show a dramatic decrease in the steady state tension compared to axons treated with a non-specific morpholino as shown in Fig. 4C. In some cases force values were so low that determination of rest tension 𝒯_0_ was not possible. The extent of knock-down was quantified using antibody labelling and the result is shown in the inset of Fig. 4C, demonstrating a correlation between reduction in spectrin content and strain-dependent steady state tension shown in the main plot. We could not observe any difference in growth characteristics or caliber between knock-down and control cells. Thus, these results clearly demonstrate the significance of the actin-spectrin skeleton in axonal response to stretch deformations. We then turned to theoretical modelling to gain further insight into how this skeleton may contribute to the observed axonal response.

### Folding-unfolding of spectrin buffers axon tension

The axonal cytoskeleton is complex (Fig. 1) [22] and delineating the contributions of the different components is not trivial. It has been generally assumed that bundled microtubules are the main mechanical element in axons [23, 24]. However, our results demonstrate that F-actin and spetcrin play a very prominent role in axonal mechanical response. Furthermore, we see that axons behave as strain-softening, viscoelastic solids at long times, unlike a fluid-like state predicted by pure microtubule based models that invoke cross-link detachments [23, 24]. For these reasons, and to obtain some insight into how the actin-spectrin skeleton contributes, we first consider the actin-spectrin skeleton alone and model its response to mechanical stretch.

When an axon is suddenly stretched at constant strain, the tension is not constant, but instead relaxes over time (Fig. 2B) till it reaches a non-zero steady-state value, indicating that dissipative processes are occurring at the microscopic scale. The observation of a steady state tension at long times precludes unbinding processes (it is also precludes unbinding of microtubules-associated or actin-associated crosslinkers), such as between spectrin tetramers and actin rings, as repeated unbinding events will allow tension to relax all the way to zero. Therefore, we model tension relaxation as arising from unfolding/re-folding of repeats along a spectrin tetramer, which is known to yield softening at the single tetramer level [16, 25]. These unfolding events lead to dissipation of stored elastic energy.

With these assumptions in mind, our model consists of *M* spectrin tetramers per axon cross-section, such that tetramers are longitudinally linked by actin filaments, as shown in Fig. 5A. Note that the model holds even if actin is not present as rings but as short filaments that interconnect the tetramers like in RBCs. A spectrin tetramer is modeled as a polymer chain with *N* = 76 repeats [25], and each repeat can be in an unfolded (*u*) or a folded (*f*) configuration. Then, we let *N*_u_(*t*) be the number of unfolded repeats on a tetramer at time *t*; therefore *N*_f_ (*t*) = *N* − *N*_u_(*t*) is the number of folded ones. Transitions from *u* ↔ *f* depend on the typical tension per spectrin tetramer, 𝒯_s_, in an axonal cross-section, that is, 𝒯_s_ ≈ 𝒯 /*M*, and, in turn, 𝒯_s_ will depend on the state of folding along the tetramer.

**FIG. 5.**
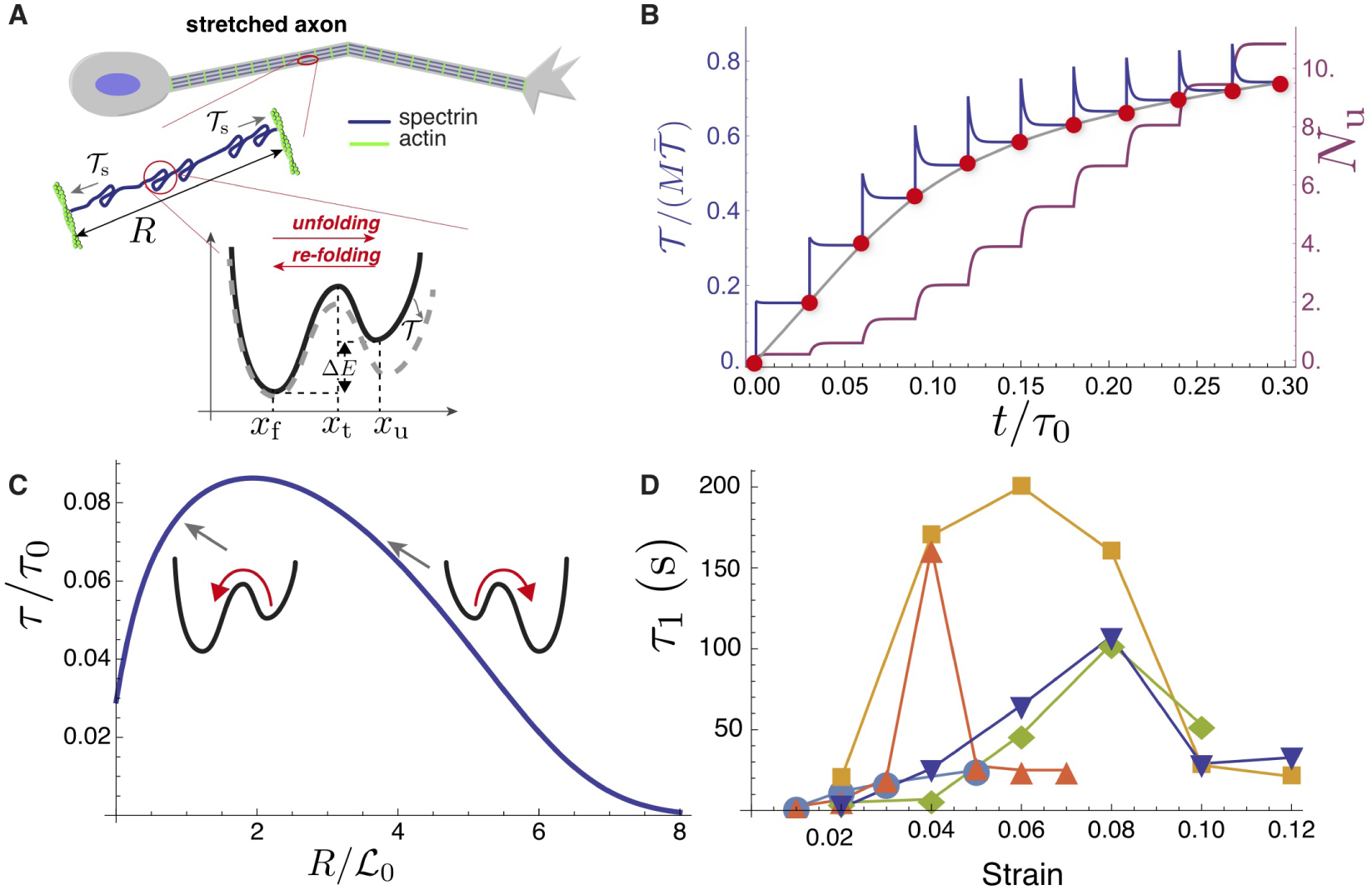
Theoretical model recapitulates strain-softening and tension relaxation. **(A)** Schematic illustration of stretched axon, showing unfolding and re-folding of spectrin repeats as underlying tension relaxation mechanism. Zoom shows schematic illustration of energy landscape for a unfolding/re-folding of a spectrin repeat. **(B)** Model calculation for a single spectrin tetramer for multiple step-strain protocol. Tension versus time (purple) shows a jump after strain increment is applied, followed by relaxation to a steady-state value (red points, passed through by the equilibrium force versus extension curve (gray)). This relaxation coincides with progressive spectrin unfolding, as indicated by the number of unfolded repeats, *N*_u_ (burgundy). Note that the tension scale used is 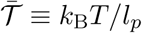. **(C)** Tension relaxation after a small change in applied strain is approximately exponential, with a relaxation time, *τ*, that depends non-monotonically on strain. The inset cartoons reveal that *τ* is largely controlled by re-folding at low strain, and by unfolding at high strain. **(D)** Fitting the tension relaxation data obtained from experiments like the one shown in Fig. 2B to a double-exponential function (Figs. S17, S18) reveals a long relaxation time with a qualitatively similar dependence on strain as with the model (Figs. S19, S20). Model calculations in B and D were done using the following parameters: *k*_B_*T* = 4 pN.nm, *l*_*p*_= 0.6 nm (average of values obtained from [16]), ℒ_0_ = 200 nm, Δℒ = 31.7 nm, *N* = 76, *x*_t_ − *x*_f_ = 3.5 nm, *x*_u_ − *x*_t_ = 3.5 nm, Δ*E* = 8 pN.nm.

In a mean-field approach, *N*_u_(*t*) evolves according to the kinetic equation

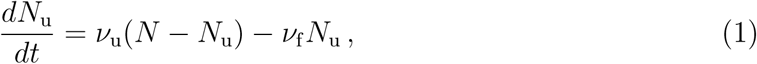

where *ν*_u_ and *ν*_f_ are, respectively, the unfolding and folding transition rates, and are assumed to depend on the tension [25, 26]:

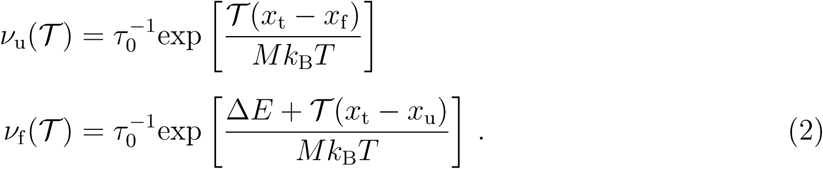

Here, the inverse time constant 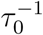 is the unfolding rate at zero tension. Furthermore, *x*_f_, *x*_t_, and *x*_u_ are position-related reaction coordinates in the folded, transition, and unfolded states, such that *x*_u_ > *x*_t_ > *x*_f_ (Fig. 5A). Finally, *k*_B_ is Boltzmann’s constant, *T* is temperature, and Δ*E* > 0 is the energy difference between the *u* and *f* states.

In the next step, we model the force-extension relation of a spectrin tetramer using an interpolation formula from the wormlike chain (WLC) model for polymer elasticity [16, 27]:

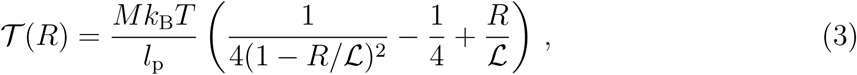

where *R* is the end-to-end extension of the tetramer; *l*_p_ is its persistence length; and ℒ is its contour length. A key ingredient in the model is that ℒ is not constant, but rather depends on the folding state of the molecule. That is,

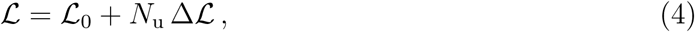

where ℒ_0_ ≈ 200 nm is the contour length with all repeats folded and Δℒ ≈ 30 nm is the gain in length when a repeat unfolds [16].

With this model in hand, we can further explore the axon tension relaxation and strain-softening seen experimentally. By solving Eqs. (1)-(4) for a sequence of equal strain steps, each applied at constant time intervals, our model can qualitatively reproduce the axon stretch response (Fig. 2B). Following a jump in strain, the tension rises quickly and then relaxes, as spectrin repeats progressively unfold (Fig. 5B). At long times after the strain step, the tension tends to a steady-state 𝒯_ss_, such that the locus of values of 𝒯_ss_ follows the steady-state tension versus strain curve (red dots in Fig. 5B). This reflects the tension buffering behavior seen in Figs. 2B, 1D(inset), Fig. S15, and Ref. [25]. This curve reveals the characteristic strain-softening effect predicted by this model: for small extension, *R*, most repeats are folded, *N*_u_ ≈ 0, and the tetramer behaves like a polymer with fixed contour length and with an elastic response 𝒯 ∝ *R*. For larger *R*, the number of unfolded repeats increases, as does the contour length and, so to speak, the reference state of the polymer. This slackening of the tetramer gives rise to softening with increasing strain (Fig. S16).

Our model also predicts a surprising behavior in the axon tension relaxation, specifically a non-monotonic dependence of the relaxation time on strain; see Fig. 5C. By subjecting each model tetramer to a sudden change in extension, from *R* to *R* + Δ*R*, and linearizing Eqs. 1-4 for small Δ*R* around the steady-state at *R*, we find exponential relaxation with time constant

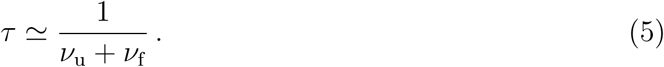

Here, *ν*_u_ and *ν*_f_ are the transition rates in the state *R* (see Suppl. Mat. Sec. 12 for details). Remarkably, this expression captures very well the experimental dependence of the tension relaxation time with strain shown in Fig. 5D, and the underlying physics can be readily understood. For small *R*, given that Δ*E*/*k*_B_*T* > 1 [25], *ν*_u_ ≪ *ν*_f_, and thus *τ* ≈ 1/*ν*_f_. That is, the *folding* rate initially dictates the rate of tension relaxation. Since, the folding rate decreases with tension, it then follows that *τ increases* with 𝒯, or equivalently with *R*. For large *R*, the energy landscape for unfolding and re-folding becomes sufficiently tilted that *ν*_u_ ≫ *ν*_f_ (Fig. 5C). Then, *τ* ≈ 1/*ν*_u_, and since *ν*_u_ increases with 𝒯, we find that the relaxation time decreases with strain.

In summary, our simple model is able to account for the main outcomes — strain-softening, non-zero steady state tension, and non-monotonic tension relaxation — of axon stretch experiments that point to a leading role played by the actin-spectrin ultrastructure.

## DISCUSSION AND CONCLUSIONS

To summarize our main results, using a custom-built, controlled-strain force apparatus, we demonstrate for the first time, to our knowledge, that (i) axons exhibit a strain-softening behavior due to their ability to buffer mechanical tension when stretched (tension homeostasis); and (ii) that their passive response behave as viscoelastic solid, with a relaxation time that depends non-monotonically on strain. We have, furthermore, unravelled hitherto unknown connections between the axon mechanical response and its cytoskeleton, by using experiments that either stabilize or de-stabilize specific elements, including the actin-spectrin skeleton. We show that apart from microtubules, (iii) F-actin and spectrin emerge as prominent contributors to the axon mechanics. As axons mature, the actin-spectrin periodic skeleton becomes more prevalent and this correlates with an increase in the Young’s moduli of the axons with age.

An order of magnitude estimate of the maximum possible contribution coming from spectrin tetramers to axonal tension can be made as follows. If we assume the width of a spectrin molecule to be 10 nm, and take a typical axon diameter as 1 µm, the maximum number of spectrin molecules (for a mature axon, say) in a cylindrical cross-section can be estimated to be about *M* ∼ 300. A more conservative estimate can be made using available data from RBCs [28], where the actin-spectrin junction is reported to be 35 nm in width (with each junction connecting to an alpha and a beta spectrin), and this gives *M* ∼ 180. If we now take the force needed to unfold a spectrin subunit from AFM experiments as *f*_s_ ≃ 30 pN [16], we can estimate the axonal tension to be of the order of 𝒯_ss_ ∼ 10 − 6 nN. This is in a reasonable range when compared to the experimentally measured plateau tension for 4-DIV axons (Fig. S15). Although this is only a rough estimate, and the exact number may depend on the details of the actin rings, it suggest that spectrins could make a significant contribution to axonal mechanics.

Our experiments and theoretical analysis suggest that spectrin can contribute to strain-softening via force-dependent unfolding and re-folding of spectrin repeats. The minimal model we present ignores possible contributions from other axonal cytoskeletal elements, which will be elaborated below. However, this actin-spectrin model predicts key experimental features like the viscoelastic solid-like response and the non-trivial dependence of the tension relaxation time, which first increases and then decreases at larger strain. These two responses are possible signatures of the role of spectrin unfolding and re-folding in tension relaxation, and argues against a few other possible mechanisms mentioned in Table-S2. For example, a tension relaxation scenario involving actin-spectrin unbinding—or any other crosslink unbinding—would lead to a relaxation time that monotonically decreases with strain and a long time fluid like response. An additional feature of the force-assisted unfolding of spectrin domains is that the tension versus strain response at steady-state exhibit an extended region where the tension is only weakly dependent on the strain (Fig. 2D-inset, S15 and S16). This can have important functional significance as the actin-spectrin skeleton can protect axons against stretch deformations by acting as a tension buffer, or “shock absorber”.

There are *in-vivo* studies that reveal the importance of spectrin in axonal mechanical stability or tension. Notably, in *C. elegans*, the axons of spectrin knock-out animals are known to snap during normal wiggling of the worm, although the exact nature of axonal deformation (buckling vs stretch) that causes damage is unclear [29]. Moreover, the use of a FRET-based spectrin tension sensor has shown that *β*-spectrin in *C. elegans* axons are held under prestress; and this is further supported by axonal retraction observed after laser ablation [30]. Still, to date, there have been no studies that quantitatively relate the actin-spectrin axon architecture to its strain response. In contrast, spectrin-mediated elastic behavior leading to strain softening has been documented in experimental and theoretical studies on Red Blood Cells (RBCs), which have a membrane associated hexagonal actin-spectrin skeleton [25, 31–33]. The current view is that RBC softening is most likely due to the unfolding of spectrin domains under force [34, 35].

### Contributions to axonal mechanics from other cytoskeletal components

As shown schematically in Fig. 1, the axonal cytoskeleton is a composite structure consisting of microtubules, neurofilaments, and cortical F-actin structures other than that present as rings. Despite the variety of perturbation experiments we report, delineating the individual contributions from these elements become challenging because of the mutual dependencies between them. For example, it has been shown that disruption of microtubules using Nocodazole causes a decay of the actin-spectrin skeleton and stabilizing the filaments using Taxol increases the occurrence of this periodic scaffold [21]. Further, perturbing one component could de-stabilize others. Hence, the effects of microtubule perturbation, for instance, may be more complex than a simple change in their numbers. Microtubule bundles alone can account for softening if their cross-linkers (MAPs) exhibit binding-unbinding dynamics, but such models give a long time fluid like response [23, 24] and also cannot account for the peak in the relaxation time vs strain plot. Thus, while microtubules may be making a direct contribution to the effective moduli we measure, microtubule elasticity alone cannot account for the entire set of experimental observations we report. Another possibility is that microtubules become disrupted under dynamic loading. In fact, microtubules were shown to be weak under dynamic stretching of axons, as they undergo depolymerization at high strain or strain rates [36]. Such catastrophic decay of microtubules under stretch, if it occurs, makes the mechanical support rendered by the actin-spectrin skeleton even more critical in ensuring axonal integrity under fast stretch. However, we note that our experiments where we first stabilize microtubules with Taxol and then disrupt F-actin with Lat-A still causes a reduction in stiffness comparable to neurons treated with Lat-A alone. This suggests that the actin-spectrin skeleton is a major contributor even in presence of stabilised microtubules. Neurofilaments, too, can play a role in axon mechanics as they form ionic crosslinks and connections to microtubules via motor proteins that transport them, although *in vitro* studies show that these networks stiffen considerably before any plasticity sets in [37, 38].

### Strain stiffening and softening in other cell types

The axonal response reported here is in sharp contrast with the strain stiffening response exhibited by many cell types [39, 40]. This stiffening has its origin in the entropic nature of the F-actin cytoskeleton and has been observed in purified systems of a variety of biopolymers [38]. It is also known that the stiffening response can transition towards a softening response at higher strains, either due to the force induced detachment of transient cross-links or due to buckling of filaments [41–43]. In contrast, the axons we have studied here show only a softening response even at the smallest of explored strain values. It has been reported that eukaryotic cells show a transient softening following a fast stretch and release protocol [44]. This is a transient effect and is not reflected in the steady state modulus of the cell and has been attributed to ATP driven processes, which make the cell behave as a soft-glassy system [45]. The same cells, when subjected to a step strain protocol to measure the steady state modulus, exhibit strain stiffening [44].

In conclusion, we have demonstrated a mechanical role for the actin-spectrin skeleton in axonal response to stretch. By combining quantitative experiments and theoretical modelling, we show that axonal strain-softening could arise from this spectrin skeleton and it allows axons to undergo significant reversible deformations by acting as molecular bellows which buffer tension. Although our modelling is restricted to the actin-spectrin skeleton, how the composite structure of the axon, with its different interdependent cytoskeletal elements, responds to stretch as a whole is an interesting area for further research and modelling. Moreover, how the actin-spectrin skeleton self-assembles and how it may dynamically reorganize under stretch on long timescales during growth will be interesting to explore. Apart from revealing the unique bio-mechanical properties, these results should also motivate the design of novel biomimetic materials which can deform at constant stress while retaining the memory of its initial state. From a medical point of view, spectrin mutations are associated with neurological disorders like spinocerebellar ataxia [46] and early infantile epileptic encephalopathies [47]. Hence, apart from direct mechanical effects, how axonal integrity may be affected by spectrin specific mutations will be an area of interest for future studies.

## MATERIALS AND METHODS

### Primary Neuronal culture

Fertilised Giriraja-2 chicken eggs were obtained from Karnataka Veterinary, Animal and Fisheries Sciences University, Bangalore, India. The eggs were incubated for 8-9 days and Dorsal Root Ganglia (DRG) were isolated and dissociated using a standard protocol. Cells were then plated on well cleaned glass coverslips with an attached cylindrical glass ring (1 cm high, 12 mm dia.) to contain the culture medium. We used L-15 medium (21083-027, Thermo Fisher Scientific) thickened with 0.006 g/ml methyl cellulose (34516, ColorconID) and supplemented with 10% Fetal Bovine Serum (10100-147, Thermo Fisher Scientific), 2% D-glucose (G6152, Sigma-Aldrich), 20 ng/ml Nerve Growth Factor NGF-7S (13290-010, Thermo Fisher Scientific), and 0.5 mg/ml Penicillin-Streptomycin-Glutamine (100×) (10378-016, Thermo Fisher Scientific). Cells were incubated at 37 °C for 48 or 96 h as per experiment. Prior to force measurement experiments, neurons were incubated for 30 min in supplemented L-15 medium lacking methyl cellulose.

### Cytoskeleton treatments

All cytoskeleton perturbing agents such as Nocodazole (M1404, Sigma-Aldrich), Latrunculin-A (L5163, Sigma-Aldrich), Jasplakinolide (J7473, Thermo Fisher Scientific), and Paclitaxel (Taxol) (T7402, Sigma-Aldrich) were dissolved in DMSO. The final DMSO concentration was kept well below 1% during experiments. The concentrations of these agents are indicated in the text describing these experiments.

### Functional knockdown of *β*-II spectrin

Primary chick DRG neurons were cultured in 400 µl of supplemented L-15 medium as described above for 2-4 h at 37 °C. Neurons were transfected by adding a pre-mixed solution of 20 µM Morpholino oligomers (MO) and 2 µM Endoporter (EP) (GeneTools, LLC) to the culture medium and incubated at 37 °C for 48 h in well humidified chambers. The transfection conditions were optimized by fluorescence microscopy using a carboxyorescein tagged control MO (5’-CCTCTTACCTCAGTTACAATTTATA-3’). All neurons (n = 46) showed diffuse fluorescence in the soma and all along the axon indicating the presence of tagged MO in the cytosol. An unlabeled translation blocking MO targeted against *β*-II spectrin (5’-GTCGCCACTGTTGTCGTCATC-3’) was used for functional knock-down studies. An unlabeled non-specific morpholino (5’-CCTCTTACCTCAGTTACAATTTATA-3’) was used as a control. To quantify the extent of *β*-II spectrin knockdown, neurons transfected with either anti-*β*-II spectrin or control MOs for 48 h were rinsed in HBSS (with *Ca*^++^ & *Mg*^++^), fixed and immunostained as described below. For force measurements on MO transfected cells, the culture medium was replaced by supplemented L-15 lacking methyl cellulose 30 min prior to the experiments.

### Immunofluorescence

Neurons were fixed using 4% (w/v) paraformaldehyde (PFA) and 0.5% (v/v) glutaraldehyde in phosphate buffered saline (PBS: 5.33 mM *KCl*, 0.44 mM *KH*_2_*PO*_4_, 4.16 mM *NaHCO*_3_, 137.93 mM *NaCl*, 0.33 mM *Na*_2_*HPO*_4_, 5.55 mM D-Glucose) for 10 min at room temperature (RT). Neurons were permeabilized with 0.2% (v/v) Triton X-100 in PBS for 5 min at RT and incubated in a blocking buffer of 3% (w/v) Bovine Serum Albumin in PBS at RT for 1 h. The fixed neurons were rinsed three times with PBS and incubated with anti-*β*-II spectrin antibody (BD Biosciences, 612563; 1:1500 dilution in blocking buffer) overnight at 4 °C. After rinsing again, the neurons were incubated with anti-mouse IgG conjugated to fluorescent Alexa Fluor 568 (A-11004, Thermo Fisher Scientific) or Alexa Fluor 488 (A-11001, Thermo Fisher Scientific) used at 1:1000 dilution in the blocking buffer for 1 h at RT. Prior to mounting for microscopic examination, the cells were rinsed again and post-fixed with 4% PFA and 0.5% glutaraldehyde in PBS for 10 min at RT. For super-resolution microscopy, the neurons cultured for various days *in vitro* (DIV) were mounted in Mowiol (10% Mowiol 4-88 in poly(vinyl alcohol), 81381, Sigma-Aldrich) and DABCO (2.5% w/v, 1,4-diazobicyclo[2.2.2]octane, D27802, Sigma-Aldrich) and kept overnight in the dark at 4 °C prior to imaging.

### Microscopy

*β*-II spectrin immunofluorescence of morpholino transfected neurons were evaluated using a confocal laser scanning microscope (Leica TCS SP8, Leica Microsystems) with a 63×, 1.4NA oil immersion objective. All imaging parameters were held constant for control and anti-*β*-II spectrin MO transfected neurons. Stimulated Emission Depletion (STED) nanoscopy was carried out using a Leica TCS SP8 STED system (Leica Microsystems). The fixed and mounted samples were imaged with the STED WHITE oil objective lens (HC PL APO 100×/1.40 OIL). CMLE JM deconvolution algorithm from Huygens Professional software (version 17.04) was used for deconvolution and processing of images.

### Image Analysis and statistics

The average fluorescence intensity was measured using the segmented line tool of ImageJ software (NIH, USA) with a specified width to trace the axon of interest and to calculate the intensity per unit area of the axon. When doing this, the background intensity per unit area was measured and subtracted from the axon intensity. Origin software (OriginLab, version 9) was used for statistical analysis and graphical representations. The periodicity of *β*-II spectrin distribution was evaluated by plotting the intensity trace along the axon using Fiji/Image J. The intensity values were analyzed using the autocorrelation (autocorr) function of Matlab (MathWorks, R2018b). For 1-D autocorrelation analysis, several 1*µ*m segments were taken from each axon. A line of 1 *µ*m was drawn with a line width on the edge of the axon and the intensity profile is obtained using Image J. For these segments, 1-D autocorrelation was calculated using Matlab. Then an average of the autocorrelation was obtained for all axons. The amplitude of autocorrelation is defined as the difference between the maxima (∼200 nm(first peak)) and minima (∼100 nm) of the 1-D autocorrelation. The average amplitude is obtained by averaging over all the axons.

All data are represented as mean ± standard deviation of mean (SE) from independent experiments, unless otherwise specified. Data shown represent the number of neurons (n) analysed. Whenever data sets are small (due to practical difficulties in obtaining sufficient sample size), individual data is shown before presenting mean and SE. To avoid errors due to axon to axon variations, trends (variation with strain or with drug treatments) are shown for the same axon whenever possible. For the quantification of spectrin knock-down, statistical significance of difference in mean fluorescence intensity across pooled data sets was tested using using non parametric two-tailed Mann Whitney U (MW - U) test with p<0.05 set as the minimum level of significance. Origin software (OriginLab,version 9) and Matlab 2018b were used for all the statistical analyses and graphical representations.

### Force measurement

Measurements were made using a modified version of an optical fiber based force apparatus we had developed earlier (S. Rao, C. Kalelkar, P A Pullarkat, Rev. Sci. Instr., vol. 84, 105107 (2013)), and is shown schematically in the main text Fig. 2A. It consists of a cylindrical glass cantilever, 10–20 µm in dia. and 5–10 mm in length, fabricated by uniformly etching the end portion of a 126 µm thick single mode optical fiber. The exact length and diameter of each cantilever is measured and its force constant determined by treating it as a perfect cantilever (S. Rao, C. Kalelkar, P A Pullarkat, Rev. Sci. Instr., vol. 84, 105107 (2013)). We use the Young’s modulus value for the optical fiber obtained by loading test cantilevers by known weights and measuring the tip deflection using a horizontal microscope. The base of the cantilever was attached to a closed-loop linear piezoelectric drive (P-841.60, Physik Instruments) which has an accuracy of 1 nm and a travel range of 90 µm. The piezo was mounted on a Zeiss AxioObserver D1 (Carl Zeiss GmbH) microscope using a joystick operated XYZ-stage (XenoWorks, Sutter Instruments). The position of the tip of the cantilever is measured with a resolution of 35 nm by focussing the laser light exiting the fiber on to a Position Sensitive Detector (PSD) (S2044, Hamamatsu). The green microscope illumination light and the red laser light were separated using appropriate filters to enable simultaneous force measurements and imaging using a CCD camera (Andor Luca R604, Andor Technology) and a 40×, 0.5NA LD-A-Plan objective.

The setup was tested using another cantilever as mock sample and this gave the expected linear elastic response. Once the cantilever is placed in the container with cells, a drop of mineral oil is added on top of the culture medium to minimize convection currents due to evaporative cooling. Axons were then pulled laterally at their mid-points by extending the piezo by a distance *D* as shown in Fig. 2A of the main article. During this process we ensure that the cantilever and the axon (except at the soma and the growth-cone) are maintained slightly away from the surface of the coverslip. The resulting cantilever deflection was calculated as *D* − *d* (see Fig. 2C of the main article). The axonal strain *γ* is calculated from the initial length of the axon *L*_0_. A feedback algorithm implemented using LabView (National Instruments, v14.0) calculates the strain steps such that there is no overshoot, and maintains the strain constant for a prescribed wait time after each step. Axons with initial length in the range of about 100–200 µm were chosen for experiments. The axonal diameter was measured in each case using phase-contrast microscopy.

There are practical difficulties in performing stretching experiments on primary neurons. The cell body is poorly adherent and can easily move or detach during mechanical perturbations, especially after treating with cytoskeleton modifying drugs. This limits both the range of strain values and the duration of measurements at each strain. For these reasons, we used successive strain steps with a strain-dependent wait time when probing the non-linear response of the axon and small amplitude cyclic strains for cytoskeleton perturbation experiments. Moreover, active contractile responses to mechanical perturbations and growth cone dynamics can get superposed with the passive force relaxation process. In order to suppress active dynamics as much as possible, we performed all experiments at room temperature (25–26 °C). Axons were chosen such that their entire length was within the field of view and the data was discarded if either the soma or the growth cone moved during measurement.

## Acknowledgements

We acknowledge Seshagiri Rao for his help in improving the setup, Jagruti Pattadkal for performing the preliminary experiments, Sukh Veer for cantilever calibration, and Serene Rose David for her help with the preparation and use of reagents. We thank the IISER-Pune imaging facility for STED nanoscopy; Deepak Nair and Siddharth Nanguneri for discussions and preliminary trials on super resolution imaging. We thank Andrea Parmeggiani for valuable discussions; N V Madhusudana, Igor Mu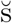evi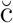, Girish Deshpande, and Patricia Bassereau for their critical reading of the manuscript. AG and PP acknowledge the Dept. of Biotechnology, Govt. of India for partial support through Grant No. BT/PR13244/GBD/27/245/2009. AG acknowledges the Science and Engineering Research Board (SERB), Govt. of India, for partial support through Grant No. EMR/2016/003730.

## Author contributions

SD, AG, ACJ & PP designed the research and analysis. Experiments and data analysis were done by SD (force measurements and imaging), SB (imaging), & NB (imaging, spectrin KD and quantification). ACJ developed the theoretical model. SD, AG, ACJ & PP interpreted data and prepared the manuscript. All authors have read and approved the final version. PP, ACJ and AG secured intramural and extramural funding.

## Competing interests

The authors declare no competing interests.

## Data availability

The raw data for plots shown in the manuscript and details of the computer codes where applicable are available from the corresponding authors on reasonable request.

## Notes

#### Summary of Updates

Both the manuscript and supplementary files have been updated. In the manuscript, texts with modified figures have been added.

## References

[1] Bruce Alberts, Alexander Johnson, Julian Lewis, Martin Raff, Keith Roberts, and Peter Walter. Molecular Biology of the Cell (Garland, New York), 2002.

[2] Ke Xu, Guisheng Zhong, and Xiaowei Zhuang. Actin, spectrin, and associated proteins form a periodic cytoskeletal structure in axons. Science, 339(6118):452–456, 2013.

[3] Elisa D’Este, Dirk Kamin, Fabian Göttfert, Ahmed El-Hady, and Stefan W Hell. STED nanoscopy reveals the ubiquity of subcortical cytoskeleton periodicity in living neurons. Cell Reports, 10(8):1246–1251, 2015.

[4] James B Phillips, Xander Smit, N De Zoysa, Andrew Afoke, and Robert A Brown. Peripheral nerves in the rat exhibit localized heterogeneity of tensile properties during limb movement. The Journal of Physiology, 557(3):879–887, 2004.

[5] Margo A Lillie, A Wayne Vogl, Kelsey N Gil, John M Gosline, and Robert E Shadwick. Two levels of waviness are necessary to package the highly extensible nerves in rorqual whales. Current Biology, 27(5):673–679, 2017.

[6] Ricardo J Andrade, Antoine Nordez, François Hug, Filiz Ates, Michel W Coppieters, Pedro Pezarat-Correia, and Sandro R Freitas. Non-invasive assessment of sciatic nerve stiffness during human ankle motion using ultrasound shear wave elastography. Journal of Biomechanics, 49(3):326–331, 2016.

[7] PV Bayly, TS Cohen, EP Leister, D Ajo, EC Leuthardt, and GM Genin. Deformation of the human brain induced by mild acceleration. Journal of Neurotrauma, 22(8):845–856, 2005.

[8] Kimberly G Harmon, Jonathan A Drezner, Matthew Gammons, Kevin M Guskiewicz, Mark Halstead, Stanley A Herring, Jeffrey S Kutcher, Andrea Pana, Margot Putukian, and William O Roberts. American medical society for sports medicine position statement: concussion in sport. British Journal of Sports Medicine, 47(1):15–26, 2013.

[9] Victoria E Johnson, William Stewart, and Douglas H Smith. Axonal pathology in traumatic brain injury. Experimental Neurology, 246:35–43, 2013.

[10] David F Meaney and Douglas H Smith. Biomechanics of concussion. Clinics in Sports Medicine, 30(1):19–31, 2011.

[11] Timothy J Dennerll, Phillip Lamoureux, Robert E Buxbaum, and Steven R Heidemann. The cytomechanics of axonal elongation and retraction. The Journal of Cell Biology, 109(6):3073–3083, 1989.

[12] Roberto Bernal, Pramod A Pullarkat, and Francisco Melo. Mechanical properties of axons. Physical Review Letters, 99(1):018301, 2007.

[13] Matthew O’Toole, Phillip Lamoureux, and Kyle E Miller. Measurement of subcellular force generation in neurons. Biophysical Journal, 108(5):1027–1037, 2015.

[14] Alireza Tofangchi, Anthony Fan, and M Taher A Saif. Mechanism of axonal contractility in embryonic drosophila motor neurons in vivo. Biophysical Journal, 111(7):1519–1527, 2016.

[15] Sampada P Mutalik, Joby Joseph, Pramod A Pullarkat, and Aurnab Ghose. Cytoskeletal mechanisms of axonal contractility. Biophysical Journal, 115(4):713–724, 2018.

[16] M Rief, J Pascual, M Saraste, and H E Gaub. Single molecule force spectroscopy of spectrin repeats: low unfolding forces in helix bundles. J Mol Biol, 286(2):553–61, Feb 1999.

[17] Nobutaka Hirokawa. Cross-linker system between neurofilaments, microtubules and membranous organelles in frog axons revealed by the quick-freeze, deep-etching method. The Journal of cell biology, 94(1):129–142, 1982.

[18] Seshagiri Rao, Chirag Kalelkar, and Pramod A. Pullarkat. Optical fiber-based force transducer for microscale samples. Review of Scientific Instruments, 84(10):105107, 2013.

[19] Hui Ouyang, Eric Nauman, and Riyi Shi. Contribution of cytoskeletal elements to the axonal mechanical properties. Journal of biological engineering, 7(1):21, 2013.

[20] Thomas Grevesse, Borna E Dabiri, Kevin Kit Parker, and Sylvain Gabriele. Opposite rheological properties of neuronal microcompartments predict axonal vulnerability in brain injury. Scientific reports, 5:9475, 2015.

[21] Guisheng Zhong, Jiang He, Ruobo Zhou, Damaris Lorenzo, Hazen P Babcock, Vann Bennett, and Xiaowei Zhuang. Developmental mechanism of the periodic membrane skeleton in axons. Elife, 3:e04581, 2014.

[22] Christophe Leterrier, Pankaj Dubey, and Subhojit Roy. The nano-architecture of the axonal cytoskeleton. Nature Reviews Neuroscience, 18(12):713, 2017.

[23] Rijk de Rooij and Ellen Kuhl. Physical biology of axonal damage. Frontiers in Cellular Neuroscience, 12:144, 2018.

[24] Hossein Ahmadzadeh, Douglas H Smith, and Vivek B Shenoy. Mechanical effects of dynamic binding between tau proteins on microtubules during axonal injury. Biophysical Journal, 109 (11):2328–2337, 2015.

[25] Qiang Zhu and Robert J Asaro. Spectrin folding versus unfolding reactions and rbc membrane stiffness. Biophysical Journal, 94(7):2529–2545, 2008.

[26] GI Bell. Models for the specific adhesion of cells to cells. Science, 200(4342):618–627, 1978.

[27] John F Marko and Eric D Siggia. Stretching DNA. Macromolecules, 28(26):8759–8770, 1995.

[28] S C Liu, L H Derick, and J Palek. Visualization of the hexagonal lattice in the erythrocyte membrane skeleton. The Journal of Cell Biology, 104(3):527–536, 1987.

[29] Marc Hammarlund, Erik M Jorgensen, and Michael J Bastiani. Axons break in animals lacking *β*-spectrin. Journal of Cell Biology, 176(3):269–275, 2007.

[30] Michael Krieg, Alexander R Dunn, and Miriam B Goodman. Mechanical control of the sense of touch by *β*-spectrin. Nature Cell Biology, 16(3):224, 2014.

[31] James CM Lee and Dennis E Discher. Deformation-enhanced fluctuations in the red cell skeleton with theoretical relations to elasticity, connectivity, and spectrin unfolding. Biophysical Journal, 81(6):3178–3192, 2001.

[32] Nir S Gov. Active elastic network: cytoskeleton of the red blood cell. Physical Review E, 75 (1):011921, 2007.

[33] Ju Li, George Lykotrafitis, Ming Dao, and Subra Suresh. Cytoskeletal dynamics of human erythrocyte. Proceedings of the National Academy of Sciences, 104(12):4937–4942, 2007.

[34] Colin P Johnson, Hsin-Yao Tang, Christine Carag, David W Speicher, and Dennis E Discher. Forced unfolding of proteins within cells. Science, 317(5838):663–666, 2007.

[35] Christine C Krieger, Xiuli An, Hsin-Yao Tang, Narla Mohandas, David W Speicher, and Dennis E Discher. Cysteine shotgun–mass spectrometry (cs-ms) reveals dynamic sequence of protein structure changes within mutant and stressed cells. Proceedings of the National Academy of Sciences, 108(20):8269, 2011.

[36] Min D Tang-Schomer, Ankur R Patel, Peter W Baas, and Douglas H Smith. Mechanical breaking of microtubules in axons during dynamic stretch injury underlies delayed elasticity, microtubule disassembly, and axon degeneration. The FASEB Journal, 24(5):1401–1410, 2010.

[37] Norman Y Yao, Chase P Broedersz, Yi-Chia Lin, Karen E Kasza, Frederick C MacKintosh, and David A Weitz. Elasticity in ionically cross-linked neurofilament networks. Biophysical journal, 98(10):2147–2153, 2010.

[38] Cornelis Storm, Jennifer J Pastore, Fred C MacKintosh, Tom C Lubensky, and Paul A Janmey. Nonlinear elasticity in biological gels. Nature, 435(7039):191, 2005.

[39] Pablo Fernández, Pramod A Pullarkat, and Albrecht Ott. A master relation defines the nonlinear viscoelasticity of single fibroblasts. Biophysical Journal, 90(10):3796–3805, 2006.

[40] Philip Kollmannsberger and Ben Fabry. Linear and nonlinear rheology of living cells. Annual Review of Materials Research, 41:75–97, 2011.

[41] ML Gardel, Jennifer Hyunjong Shin, FC MacKintosh, L Mahadevan, P Matsudaira, and DA Weitz. Elastic behavior of cross-linked and bundled actin networks. Science, 304(5675): 1301–1305, 2004.

[42] Ovijit Chaudhuri, Sapun H Parekh, and Daniel A Fletcher. Reversible stress softening of actin networks. Nature, 445(7125):295, 2007.

[43] Jan A Åström, PB Sunil Kumar, Ilpo Vattulainen, and Mikko Karttunen. Strain hardening, avalanches, and strain softening in dense cross-linked actin networks. Physical Review E, 77 (5):051913, 2008.

[44] Xavier Trepat, Linhong Deng, Steven S An, Daniel Navajas, Daniel J Tschumperlin, William T Gerthoffer, James P Butler, and Jeffrey J Fredberg. Universal physical responses to stretch in the living cell. Nature, 447:592–595, 2007.

[45] Xavier Trepat, Guillaume Lenormand, and Jeffrey J Fredberg. Universality in cell mechanics. Soft Matter, 4(9):1750–1759, 2008.

[46] Yoshio Ikeda, Katherine A Dick, Marcy R Weatherspoon, Dan Gincel, Karen R Armbrust, Joline C Dalton, Giovanni Stevanin, Alexandra Dürr, Christine Zühlke, Katrin Bürk, et al. Spectrin mutations cause spinocerebellar ataxia type 5. Nature Genetics, 38(2):184, 2006.

[47] Yu Wang, Tuo Ji, Andrew D Nelson, Katarzyna Glanowska, Geoffrey G Murphy, Paul M Jenkins, and Jack M Parent. Critical roles of *α*ii spectrin in brain development and epileptic encephalopathy. The Journal of Clinical Investigation, 128(2), 2018.

