## Supplementary Material for "The axonal actin-spectrin lattice acts as a tension buffering shock absorber"

### *Laboratoire Matière et Systèmes Complexes,*
*Université Paris-Diderot, 75205 Paris Cedex, France*

#### I. PHOTOGRAPH OF THE FORCE DEVICE

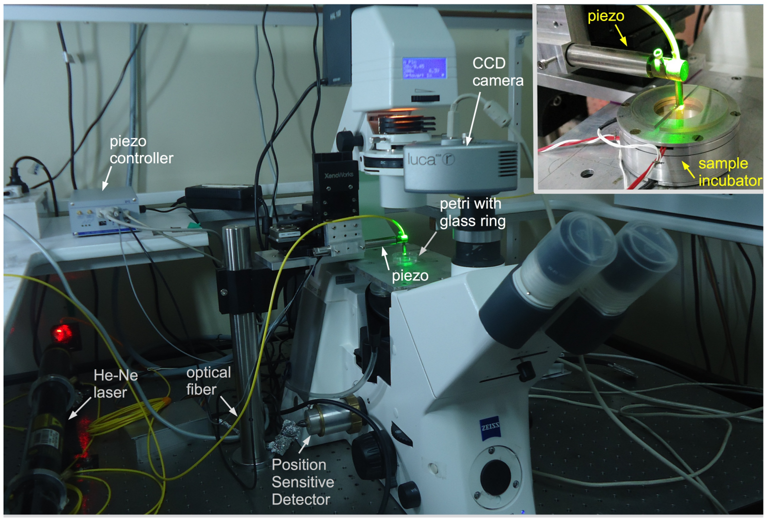


Fig. S1. Photograph of the home developed optical fiber based force apparatus used to investigate the mechanical properties of axons. The piezo drive and the Position Sensitive Detectors are interfaced to a computer to operate the setup in a strain control mode where a feedback loop maintains a prescribed strain value. More details are given in the main text. The inset shows the sample incubator. The etched glass cantilever is not visible in the image but is schematically shown in Fig 1A of the main text and the tip can be seen in the microscope images in Fig. 1B.

#### II. LONG TIME BEHAVIOR OF STRETCHED AXONS

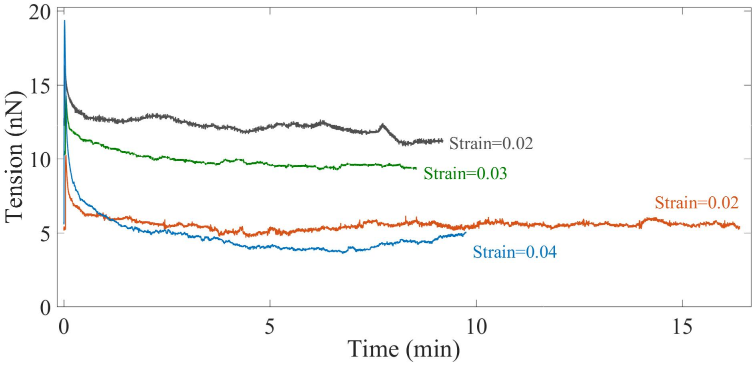


Fig. S2. Long time axonal response after the application of a single strain step shows that the force relaxes to a steady state value. This shows that axons behave as viscoelastic solid tubes at these timescales.

#### III. DETERMINATION OF REST TENSION $\mathcal{T}_0$

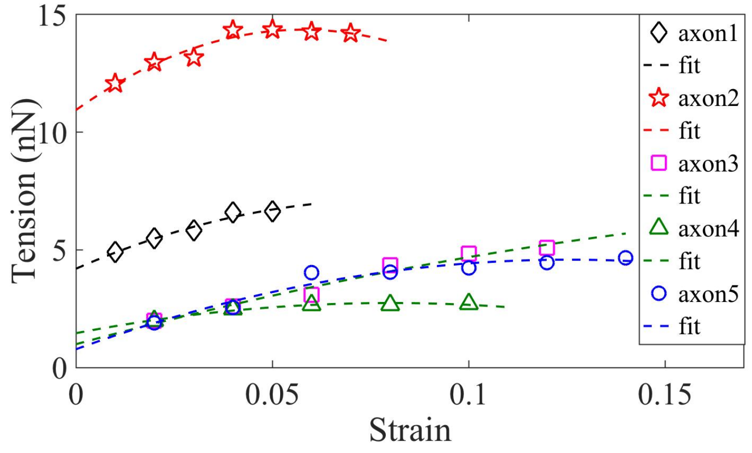


Fig. S3. Axons are under pre-stress (rest tension) even before the application of any strain. This rest tension can be obtained by extrapolating the tension vs strain plot to zero strain. This is done by fitting the data to an equation of the form $-ax^2 + bx + c$, which is a simplified form of the tension expression derived from the model presented in the main text.

#### IV. NORMALIZED YOUNG'S MODULI

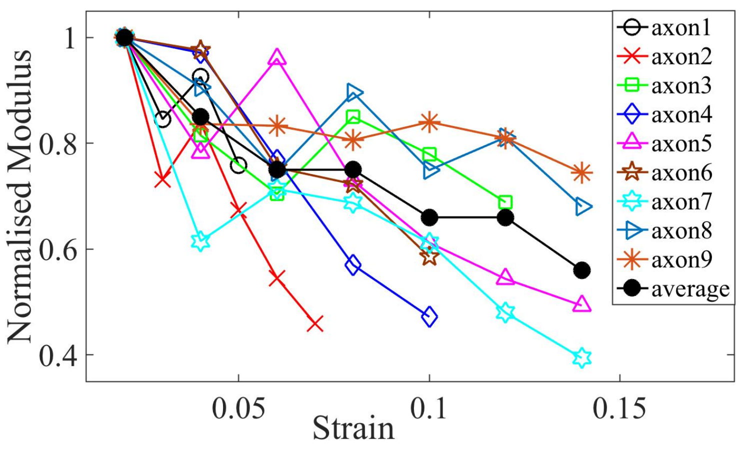


Fig. S4. Elastic moduli for different axons of age 2-DIV normalised by the first strain value which is at 0.02. The plots show softening of modulus with increasing strain. The average is also shown.

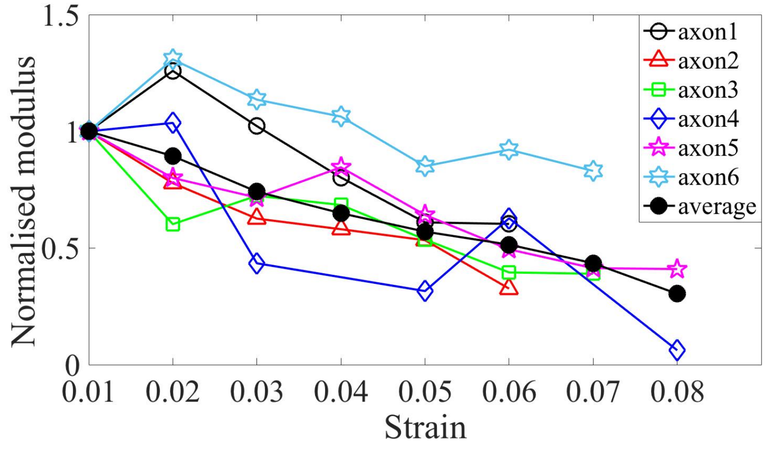


Fig. S5. Elastic moduli for different axons treated with Taxol, normalised by the first strain value which is at 0.01, which indicates softening in modulus with increasing strain. The average is also shown.

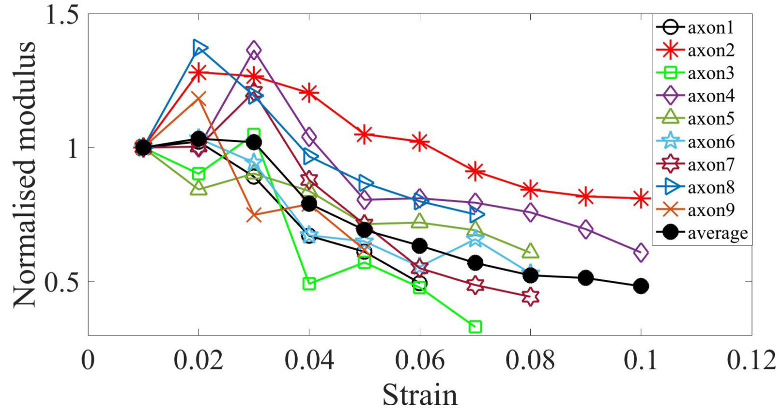


Fig. S6. Elastic moduli for different axons treated with Jasplakinolide, normalised by the first strain value which is at 0.01, which indicates softening in modulus with increasing strain. The average is also shown.

#### V. CYCLIC STRAIN PROTOCOL

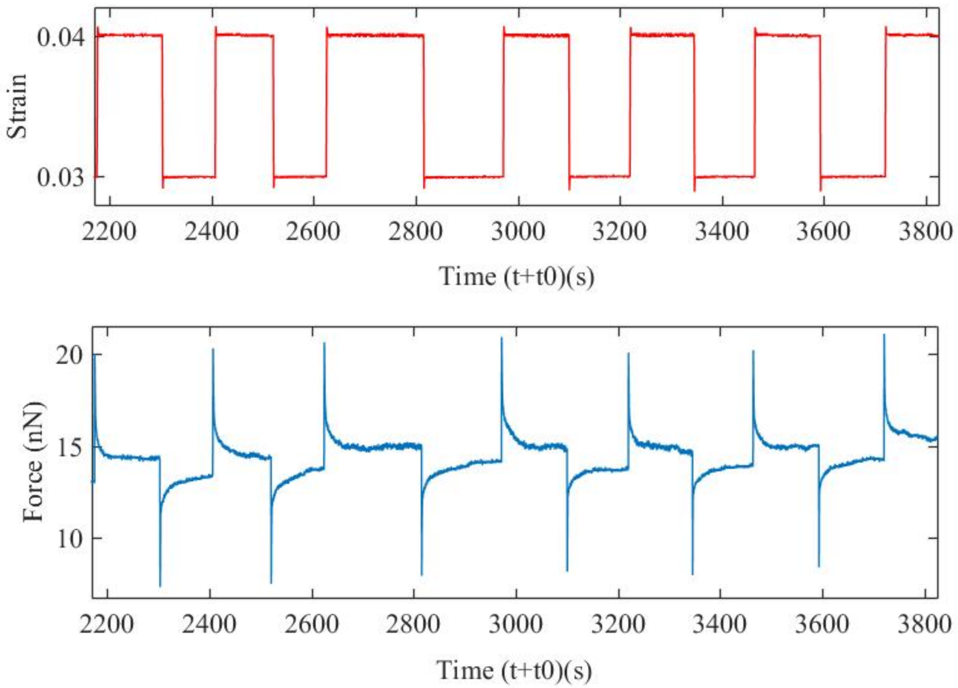


Fig. S7. An example of an axon that was pulled to a strain of 3% and then cyclic strain has been applied over an extended period to show that there is no damage with number of cycles. The data also show that the force evolution for up and down steps are similar except for sign. This protocol is then used to study the effect of specific cytoskeleton disrupting agents–Latrunculin-A for F-actin, Nocodazole for microtubules and anti-$\beta$-II spectrin morpholino for spectrin.

#### VI. FORCE RELAXATION OF DRUG TREATED CELLS

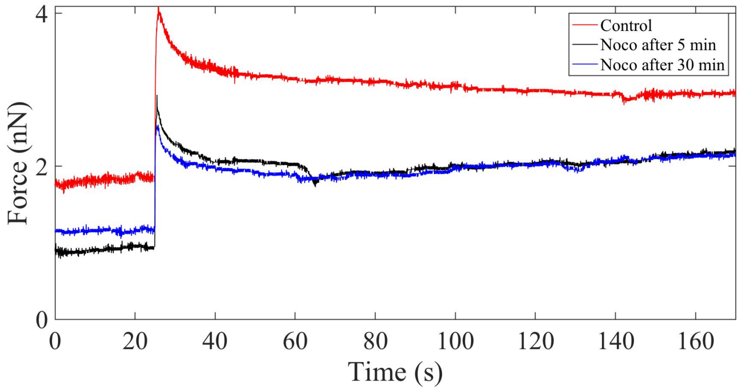


Fig. S8. Plot showing the force relaxation before and after application of 10 $\mu$M Nocodazole (noco) for 5 min and 30 min. There is no appreciable change in the relaxation time (at 2% strain) after microtubule disruption.

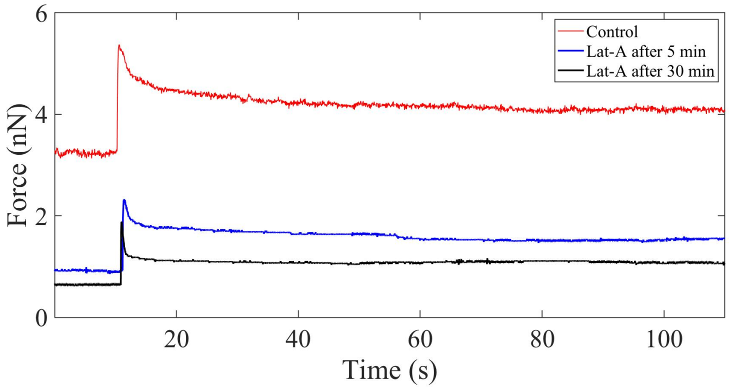


Fig. S9. Plot showing the force relaxation before and after application of 1 $\mu$M Latrunculin-A (Lat-A) for 5 min and 30 min. Unlike in the case of microtubule disruption, F-actin depolymerization causes a significant reduction in the modulus (at 2% strain) and the relaxation time.

#### VII. QUANTIFICATION OF ACTIN-SPECTRIN LATTICE

##### A. Spectrin distribution in axons (Confocal image)

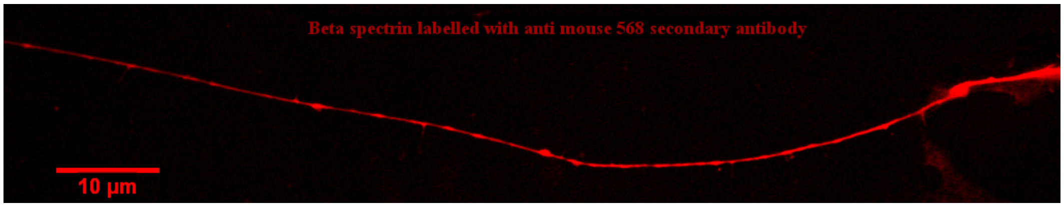


Fig. S10. Confocal image showing spectrin distribution in an axon labelled using a primary antibody anti $\beta$-II spectrin and a secondary antibody anti-mouse IgG conjugated to Alexa Fluor 568. Spectrin is seen distributed all along all axons of all ages in confocal images. The fine structure has been quantified using Super Resolution as detailed below.

##### B. Spectrin distribution in axons (STED images)

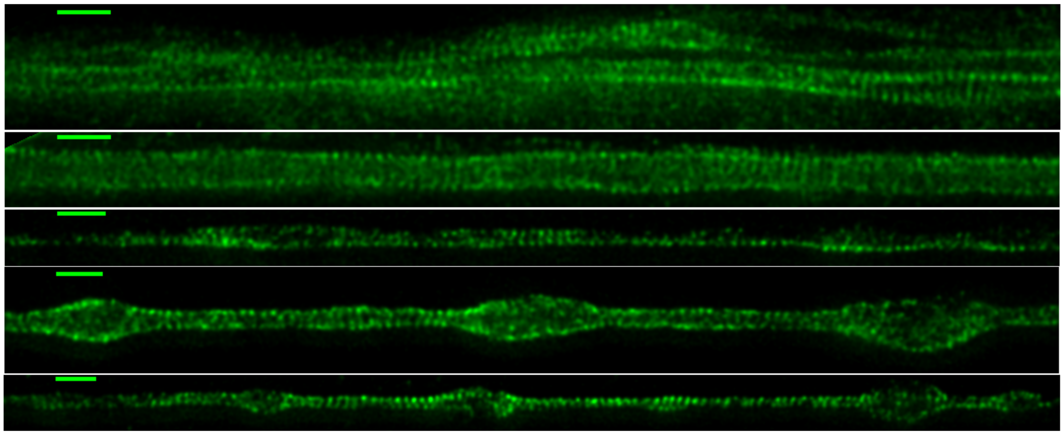

Fig. S11. Super-resolution images of 2-DIV DMSO vehicle control axons labelled using primary antibody anti$\beta$-II spectrin and secondary antibody anti-mouse IgG conjugated to Alexa Fluor 488. The rings are seen extensively (Scale bar = 1 $\mu$m). Some of the disordered patches may be due to fixation process (the presence of beading or bulges which are not seen before fixation suggest this).

##### C. STED Super Resolution Microscopy analysis of different DIV axons

| DIV | No. of axons | regular ladder | | ladder in patches | | no visible ladder | |
|---|---|---|---|---|---|---|---|
| | | Number | % | Number | % | Number | % |
| 1 | 14 | 3 | 21.4 | 3 | 21.4 | 8 | 57.1 |
| 2 | 13 | 4 | 30.8 | 6 | 46.2 | 3 | 23.1 |
| 3 | 13 | 8 | 61.5 | 5 | 38.5 | 0 | 0 |
| 4 | 10 | 7 | 70.0 | 3 | 30.0 | 0 | 0 |
| 5 | 13 | 10 | 76.9 | 3 | 23.1 | 0 | 0 |

TABLE S1. STED image analysis for quantification of rings: Here we quantify the occurrence of periodic rings (ladders) seen at various DIVs within an imaging field of view. It is evident from the table that as axon mature the rings develop all over the axons.

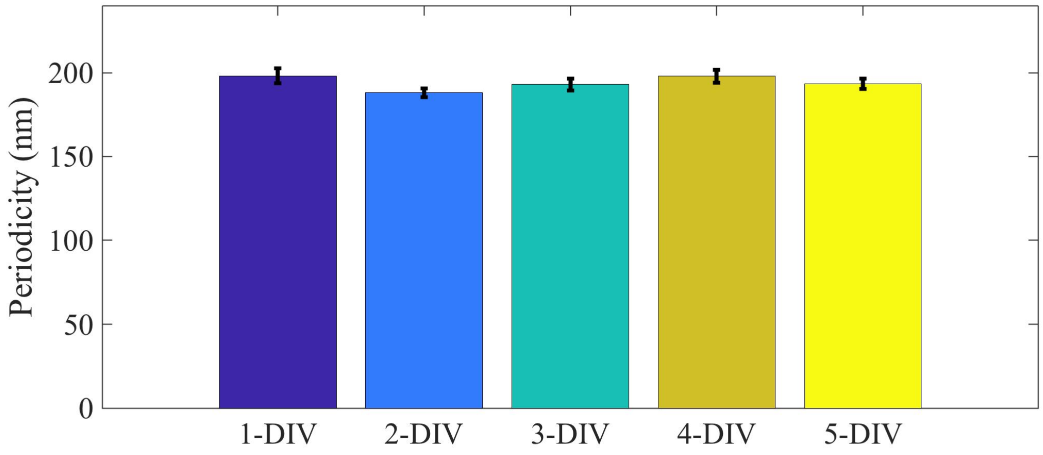


Fig. S12. Periodicity of spectrin lattice in axons of different age measured using STED microscopy. Number of axons used in each case was n(1-DIV) = 23, n(2-DIV) = 23, n(3-DIV) = 24, n(4-DIV) = 20, n(5-DIV) = 24 This analysis was done using LAS AF version 3.3.0.10134 from Lieca Micro-systems by drawing a line along the axon and the average distance between adjacent peaks were measured for each axon.

##### D. STED Super Resolution Microscopy analysis of drug treated axons

We did super-resolution imaging of 2-DIV cells after treating them with the stabilizing drugs Taxol or Jasplakinolide for 30 min. The periodic skeleton is well preserved after treatment with either of these drugs. In vehicle control, 32 out of 36 showed rings; after treatment with Jasplakinolide, 28 out of 31 showed rings; and after exposure to Taxol, 31 out of 32 showed rings. We also did treatment with Latrunculin-A and no clear signature of periodic rings could be seen in 17 out of 19 axons.

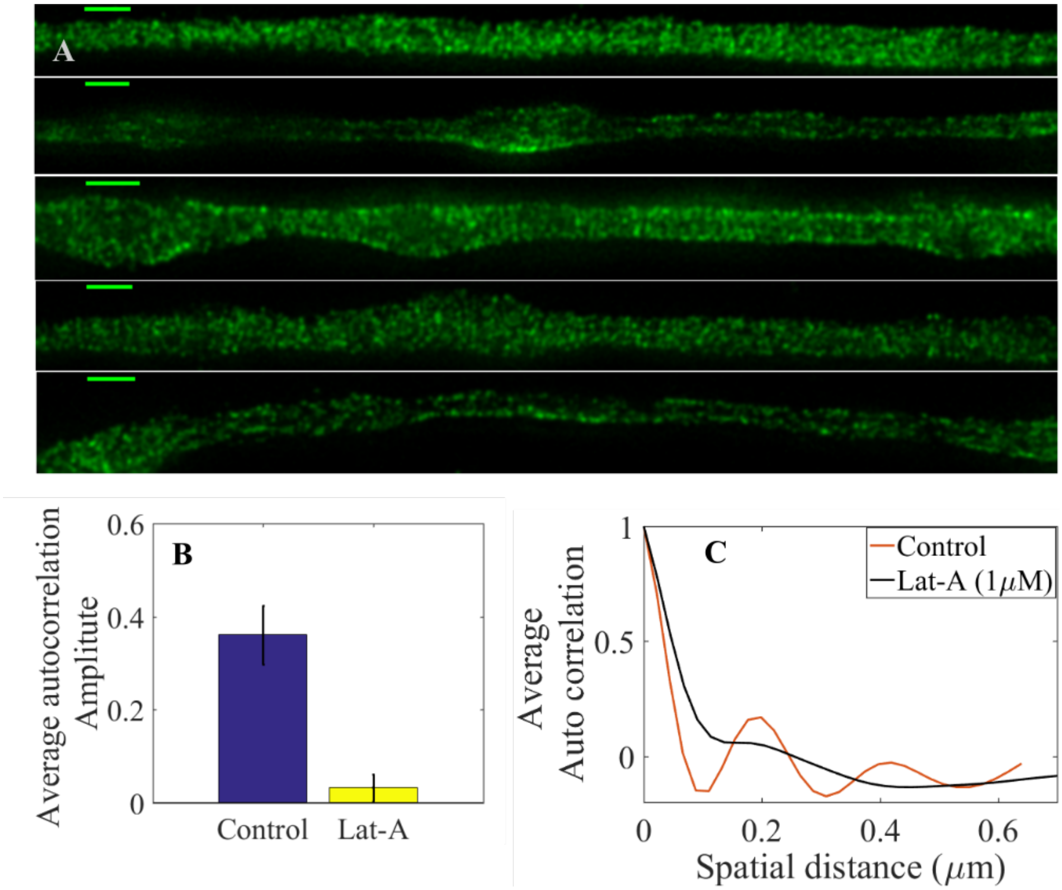


Fig. S13. (A) Super-resolution images of 2-DIV axons treated for 30 min with 1 $\mu$M Lat-A labelled using primary antibody anti$\beta$-II spectrin and secondary antibody anti-mouse IgG conjugated to Alexa Fluor 488 (Scale bar = 1 $\mu$m). It is clearly seen that the periodicity seen in control axons (Fig. S11) is absent in these images. (B) 1-D autocorrelation amplitude for several axon segments of 1$\mu$m averaged over many axons (n=10) which suggests that the spectrin periodicity is disrupted in Lat-A treated axons. (C) Intensity-intensity autocorrelation of DMSO vehicle control axons and those treated with Lat-A (1$\mu$M) for 30 min, averaged over many axons (n=10). Vehicle control axons show a clear periodicity of ∼200 nm and strong correlation where as Lat-A treated axons show very weak correlation (see Material and Methods for details).

#### VIII. REST TENSION VARIES WITH DAYS IN CULTURE (DIV)

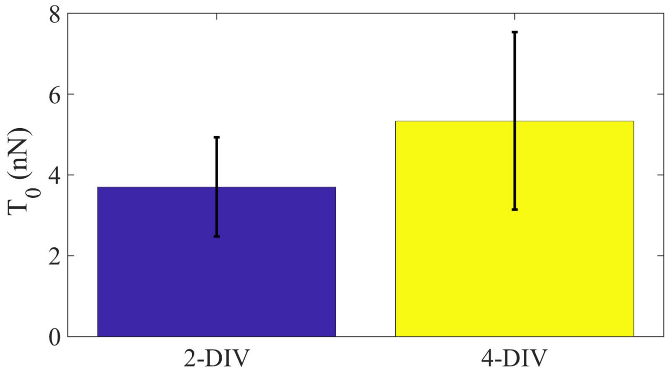


Fig. S14. Plot showing the difference in rest tension $\mathcal{T}_0$ with age for 2-DIV and 4-DIV axons. 2-DIV: n = 10, mean = 3.7, SE = 1.2; 4-DIV: n = 7, mean = 5.3, SE = 2.2. Note that the actin-spectrin lattice becomes more prominent and extensive with DIV.

#### IX. STEADY STATE TENSION VS. STRAIN IN 4-DIV AXONS

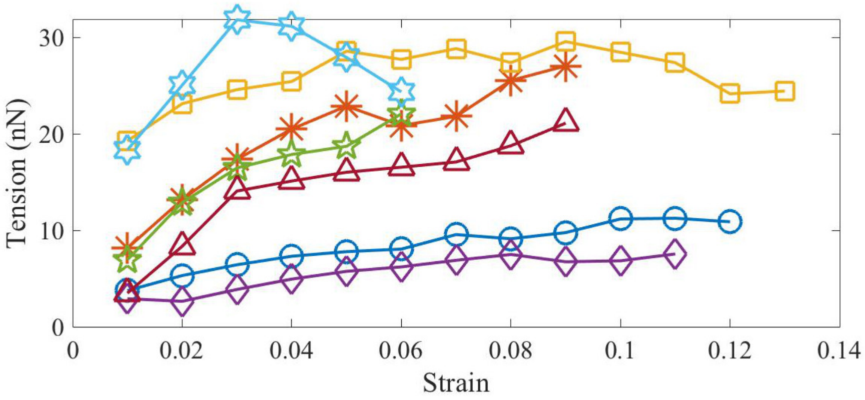


Fig. S15. Evolution of the steady state tension as a function of strain for axons grown for four days in culture (4-DIV).

#### X. ANALYSIS OF TENSION RELAXATION

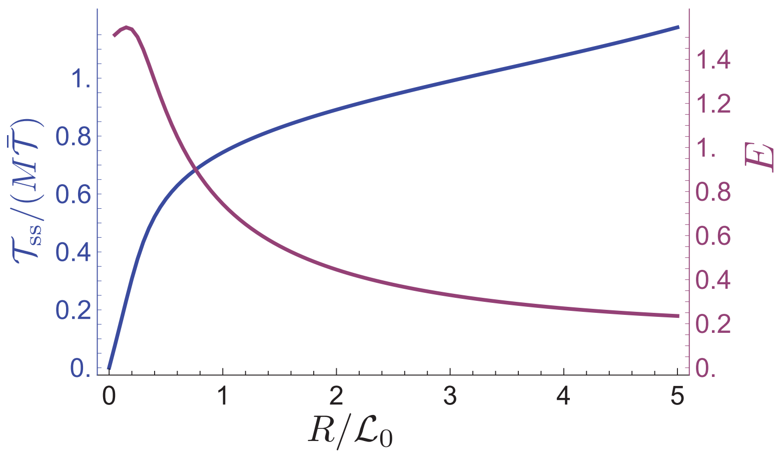


Fig. S16. Equilibrium tension and Young's modulus versus extension curve for a single spectrin tetramer. The tension curve (blue) is obtained by solving Eqs. 1-4 of the main text at equilibrium ($dN_u/dt = 0$). The tension is normalized by the number of spectrin molecules per cross-section, $M$, and by the tension scale $\bar{\mathcal{T}} \equiv k_B T/l_p$. The Young's modulus curve (purple) is obtained by dividing the normalized tension by the spectrin end-to-end extension $R/\mathcal{L}_0$. The small bump in the $E$ vs. extension curve is related to the entropic elastic contribution in the wormlike chain interpolation formula (first term on the right in Eq. 3 of the main text). Since axons are under tension at rest, with a distance of $R \approx 200$ nm between adjacent actin rings, this bump is not seen experimentally. The parameters used for both plots are: $k_B T = 4$ pN.nm, $l_p = 0.6$ nm, $\mathcal{L}_0 = 200$ nm, $\Delta\mathcal{L} = 31.7$ nm, $M = 76$, $x_t - x_f = 3.5$ nm, $x_u - x_t = 3.5$ nm, $\Delta E = 8$ pN.nm.

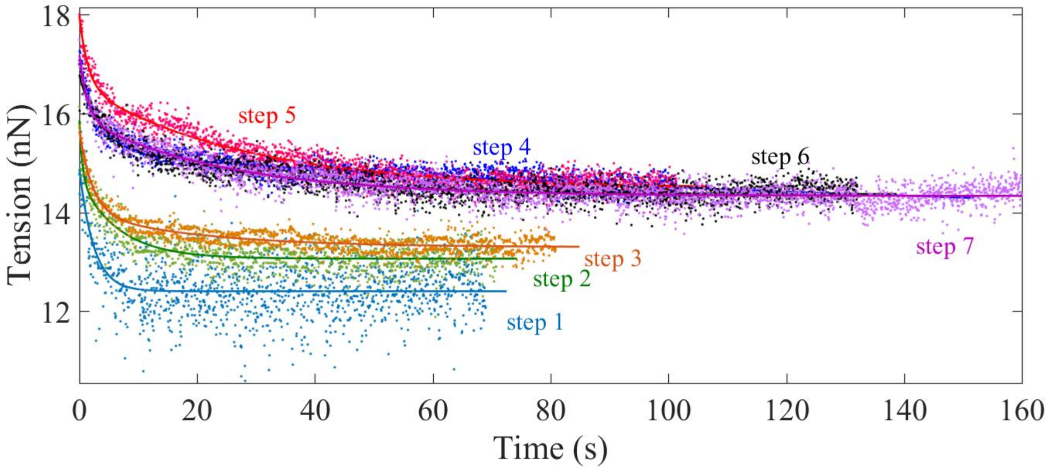


Fig. S17. Plots showing double exponential fits to the tension relaxation data from multi-step experiment (2-DIV axons) using $\mathcal{T}(t) = A\exp(-t/\tau_1) + B\exp(-t/\tau_2) + C)$ where A, B, and C are positive constants and $\tau_1$ and $\tau_2$ are the two relaxation times.

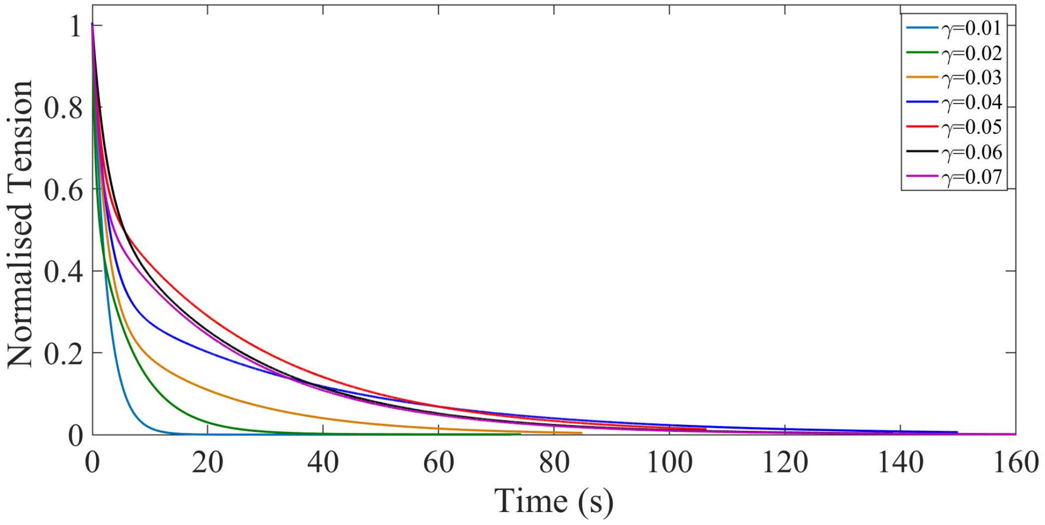


Fig. S18. The fitted curves for tension relaxation after each step were normalized by the tension immediately after each step and plotted together to better illustrate the variation in relaxation time with step number. The normalized tension is calculated as $\mathcal{T}_N = [\mathcal{T}(t) - \mathcal{T}_{\rm ss}]/(\mathcal{T}_{\rm peak} - \mathcal{T}_{\rm ss})$, where $\mathcal{T}(t)$ is the axonal tension, $\mathcal{T}_{\rm peak}$ is the peak value of tension for each step and $\mathcal{T}_{\rm ss}$ is the steady state tension after the force has relaxed.

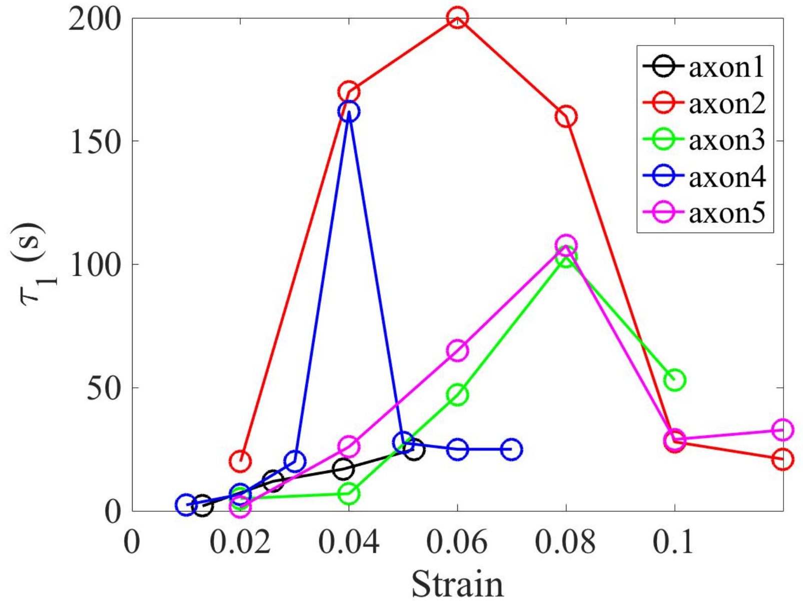


Fig. S19. Plot showing the long relaxation time as a function of strain for different 2-DIV axons.

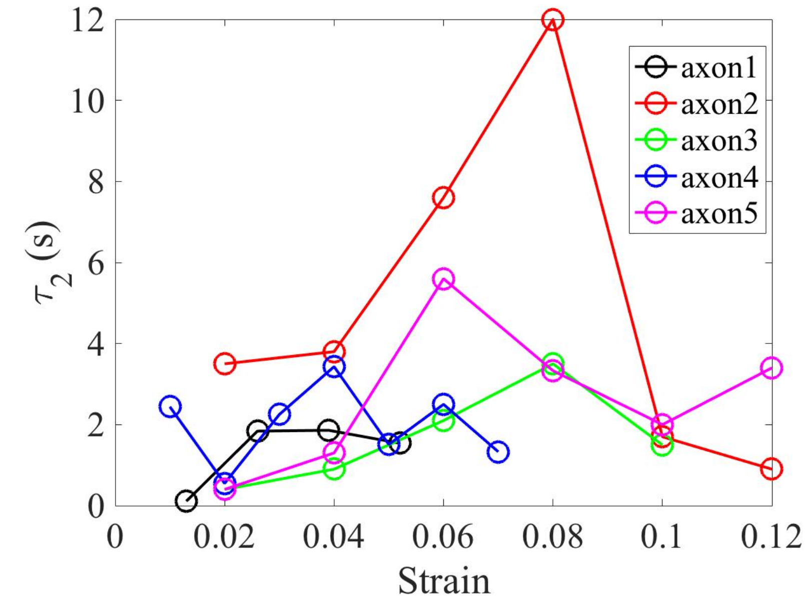


Fig. S20. Plot showing the short relaxation time as a function of strain for different 2-DIV axons.

#### XI. COMMENT ON DEPENDENCE OF TENSION RELAXATION TIME ON STRAIN

In this section, we briefly justify Eq. 5, which accounts for the non-monotonic dependence of the relaxation time on strain. To see this, starting at steady-state the model axon, as described in the main text, is subjected, at some $t = 0$, to a sudden, but small (constant) change in extension, from $R$ to $R + \Delta R$. Following this change in strain, $\mathcal{T}$ and $N_{\mathrm{u}}$ change from their steady state values, $\mathcal{T}_{\mathrm{ss}}$ and $N_{\mathrm{u,ss}}$, at extension $R$ by small amounts $\Delta\mathcal{T}$ and $\Delta N_{\mathrm{u}}$. Right after the change in strain, there is an elastic increase in tension, followed by relaxation, as spectrin repeats gradually unfold; see Fig. 4B. By linearizing Eqs. 1–4 around the steady-state at $R$, we find an exponential relaxation for $\Delta\mathcal{T}$ and for $\Delta N_{\mathrm{u}}$, with inverse time constant

$$\tau^{-1} = (\nu_{\mathrm{u}} + \nu_{\mathrm{f}}) + \left.\frac{\partial\mathcal{T}}{\partial N_{\mathrm{u}}}\right|_R \left(\frac{-\nu_{\mathrm{u}}(N - N_{\mathrm{u,ss}})(x_{\mathrm{t}} - x_{\mathrm{f}}) + \nu_{\mathrm{f}} N_{\mathrm{u,ss}}(x_{\mathrm{t}} - x_{\mathrm{u}})}{M k_{\mathrm{B}} T}\right), \qquad \text{(S 1)}$$

where the transition rates $\nu_{\mathrm{u}}$ and $\nu_{\mathrm{f}}$ and the partial derivative of $\mathcal{T}$ with respect to $N_{\mathrm{u}}$ are all evaluated at the steady state with extension $R$. We can obtain a simpler expression for the relaxation time, which then allows us to understand its non-monotonic dependence on strain, as seen experimentally. Noting that $\partial\mathcal{T}/\partial N_{\mathrm{u}} \sim \mathcal{T}\Delta\mathcal{L}/\mathcal{L} \sim 0.03\,\mathcal{T}$ for $N_{\mathrm{u}} \sim 30$, we can show numerically using Mathematica (Mathematica 11.0. Wolfram Research, Inc., Champain, Illinois, 2018) that the second term in Eq. (S 1) is small compared with the first. Thus, we find that, to a very good approximation, $\tau$ can be calculated using Eq. 5.

#### XII. CROSSLINK-DETACHMENT VS DOMAIN UNFOLDING

| | Model | Strain-softening of steady state moduli | Solid-like steady state | Peak in relaxation time vs strain |
|---|---|---|---|---|
| **i** | Randomly crosslinked semiflexible polymer gel (Ref: 1 below) | Initial entropic stiffening and subsequent softening due to force dependent crosslink detachment. Crosslink unbinding causes energy dissipation and stress relaxation. | Solid-like in stiffening regime and fluid-like in softening regime. | No |
| **ii** | Aligned microtubules with force dependent crosslinks (Refs: 2,3 below) | Can exhibit direct softening due to crosslink detachment as entropic effect is minimal. | Fluid-like at long times unless one invokes another parallel structure with permanent crosslinks. | No |
| **iii** | Force dependent unfolding and force dependent refolding of spectrin domains. | Exhibits direct strain softening when spectrin is in fully extended configuration (as suggested by 190 nm tetramer spacing and FRET data) Dissipation and stress relaxation due to unfolding of domains. | Solid-like at long times as unfolded domains can take load unlike detached crosslinks. | Peak in relaxation time vs strain because unfolding rate increases with force while refolding rate decreases with force. |

TABLE S2. A summary of different scenario and their expected mechanical response.

1. C.P. Broedersz and F.C. MacKintosh, *Rev. Mod. Phys., v86, p995, (2014)*
2. H. Ahmadzadeh, D.H.Smith, and V.B.Shenoy, *Biophys.J., v109, p2328, (2015)*
3. Rijk de Rooij and Ellen Kuhl, *Frontiers in Cellular Neuroscience, v12(144), (2018)*

#### XIII. DESCRIPTION OF VIDEO CLIPS

**Movie-S1:** Video showing an axon being stretched using the sequential step-strain protocol. After each step the strain is held constant via a feedback loop.
**Movie-S2:** Video recording of an axon being released from the cantilever from a pre-stretched state. Note that the axon recovers its initial length within seconds suggesting that no plastic deformation has occurred.
